# Engineering anti-amyloid antibodies with transferrin receptor targeting improves brain biodistribution and mitigates ARIA

**DOI:** 10.1101/2024.07.26.604664

**Authors:** Michelle E. Pizzo, Edward D. Plowey, Nathalie Khoury, Wanda Kwan, Jordan Abettan, Sarah L. DeVos, Claire B. Discenza, Timothy Earr, David Joy, Ming Lye-Barthel, Elysia Roche, Darren Chan, Jason C. Dugas, Kapil Gadkar, René Meisner, Jennifer Sebalusky, Ana Claudia Silva Amaral, Isabel Becerra, Roni Chau, Johann Chow, Allisa J. Clemens, Mark S. Dennis, Joseph Duque, Laura Fusaro, Jennifer A. Getz, Mihalis S. Kariolis, Do Jin Kim, Amy Wing-Sze Leung, Hoang N. Nguyen, Elliot R. Thomsen, Pascal E. Sanchez, Lu Shan, Adam P. Silverman, Hilda Solanoy, Raymond Tong, Meredith E. Calvert, Ryan J. Watts, Robert G. Thorne, Paul H. Weinreb, Dominic M. Walsh, Joseph Lewcock, Thierry Bussiere, Y. Joy Yu Zuchero

## Abstract

Although the first generation of immunotherapies for Alzheimer’s disease (AD) are now clinically approved, amyloid-related imaging abnormalities (ARIA) remain a major safety problem for this class of drugs. Here, we report an antibody transport vehicle (ATV) targeting the transferrin receptor (TfR) for brain delivery of amyloid beta (Aβ) antibodies that significantly reduced ARIA-like lesions and improved plaque target engagement in a mouse model of amyloid deposition. Asymmetrical Fc mutations (ATV^cisLALA^) allowed the molecule to selectively retain effector function only when bound to Aβ while mitigating TfR-related hematology liabilities. Mice treated with ATV^cisLALA^:Aβ exhibited broad brain parenchymal antibody distribution; in contrast, anti-Aβ IgG was highly enriched at arterial perivascular spaces where vascular Aβ localizes and likely plays a role in induction of ARIA. Importantly, ATV^cisLALA^: Aβ almost completely eliminated ARIA-like lesions and vascular inflammation associated with anti-Aβ treatment. Taken together, ATV^cisLALA^ has the potential to significantly improve both safety and efficacy of Aβ immunotherapy through enhanced biodistribution mediated by transport across the blood-brain barrier.

## Introduction

Biotherapeutics such as monoclonal antibodies (mAbs) are typically unable to cross the blood-brain barrier (BBB) in appreciable amounts following systemic dosing, limiting their use for the treatment of neurological disease. Systemically administered macromolecules, like endogenous plasma proteins, appear to enter the CNS by first reaching the CSF compartment in a size-dependent manner (Davson & Segal, 1995; Rapoport & Pettigrew, 1979). Once in the CSF, transport of macromolecules into CNS tissue is further constrained by limited diffusion at brain-CSF interfaces and variable distribution to deeper brain via perivascular spaces (Pizzo et al., 2018; Wolak et al., 2015). CNS access for systemic macromolecules is therefore a major challenge and often results in uneven distribution between superficial brain regions that display higher concentrations and deep brain regions with significantly lower concentrations. Insufficient CNS delivery has been a major motivation for the development of new approaches, particularly those aimed at enabling transport across the BBB (Lange et al., 2022; Terstappen et al., 2021).

The brain is highly vascularized and leverages a large surface area for exchange between the blood and brain as a highly efficient path to achieve improved biodistribution. Decades of research has demonstrated that binding to specific proteins expressed on brain endothelial cells can enable receptor-mediated endocytosis and/or transcytosis, with the greatest amount of work focused upon the transferrin receptor (TfR) (Bien-Ly et al., 2014; Edavettal et al., 2022; Niewoehner et al., 2014; Weber et al., 2018; Yu et al., 2011, 2014). TfR is highly expressed on brain endothelial cells (Jefferies et al., 1984) and undergoes constitutive internalization from the cell surface (Hopkins et al., 1985), resulting in a relatively high potential capacity for facilitating blood-to-brain transport. We previously described a transport vehicle (TV) where monovalent TfR binding was engineered into the human IgG1 Fc domain yielding a modular platform that does not require appended sequences and can be paired with antibodies (ATV) (Kariolis et al., 2020; Khoury et al., 2024; van Lengerich et al., 2023), enzymes (ETV) (Arguello et al., 2022; Ullman et al., 2020), other proteins (PTV) (Logan et al., 2021), or oligonucleotides (OTV) (Barker et al., 2023).

Historically, it has been challenging for TfR-binding antibodies to safely preserve effector function due to depletion of TfR-expressing erythrocyte precursor cells, such as reticulocytes, that are present in the blood and bone marrow (Castellanos et al., 2020; Couch et al., 2013; Pardridge et al., 2018). There are mixed reports as to whether this liability may be molecule architecture-dependent (Edavettal et al., 2022; Weber et al., 2018) and little data exists to directly demonstrate that architecture alone is sufficient to alleviate reticulocyte loss and subsequent anemia. As a result, TfR-based delivery platforms often introduce mutations in the IgG Fc region to prevent immune cell engagement, rendering the molecules effectorless (Edavettal et al., 2022; Kariolis et al., 2020; Logan et al., 2021; Ullman et al., 2020; van Lengerich et al., 2023). While such an approach may be suitable for many CNS targets, engagement with FcɣR may be required for efficacy for other therapeutics.

Amyloid beta (Aβ) is one such target where clinical data from multiple antibodies demonstrates that both binding to aggregated Aβ species and effector function is required for efficacy. Accumulation of Aβ plaques in brain is one of the pathological hallmarks of Alzheimer’s disease (AD; (Haass & Selkoe, 2007; Petersen et al., 2021; U.S. Food and Drug Administration Center for Biologics, 2024), and efficacy of anti-Aβ therapeutics is strongly linked to removal of plaques (Karran & De Strooper, 2022) presumably by microglia-mediated phagocytosis in an effector-function dependent manner (Bard et al., 2000; Citron, 2010; Koenigsknecht-Talboo et al., 2008). The recent clinical successes of multiple Aβ antibodies have helped solidify the amyloid hypothesis as a promising disease-modifying therapeutic approach (Glenner & Wong, 1984; Selkoe & Hardy, 2016). However, based on the expected inefficiency of brain exposure for antibodies due to their limited passage across the BBB (St-Amour et al., 2013) and physiological transport limitations once in the CSF (Abbott et al., 2018; Hladky & Barrand, 2022; Pizzo et al., 2018; Wolak et al., 2015), there is likely significant opportunity for improvement in both exposure and distribution within the brain to further enhance target engagement and efficacy. Additionally, amyloid-related imaging abnormalities (ARIA) remain as the predominant safety concern for this class of therapeutics. Although the mechanisms underlying lesions such as vascular hemorrhages and edema associated with ARIA remain incompletely understood, the occurrence of these adverse events appears to be linked to vascular amyloid deposition (i.e., cerebral amyloid angiopathy, CAA), presence of the *APOE ε4* allele, and dose level (Hampel et al., 2023; Salloway et al., 2014; R. Sperling et al., 2012; R. A. Sperling et al., 2011). These adverse events are dose-limiting and restrict their utility in certain patient populations. Strategies to further improve both the safety and efficacy of this promising class of therapeutics will therefore be critically important to their broader adoption.

Here, we describe the development and characterization ATV^cisLALA^, a TfR-based antibody transport vehicle platform paired with anti-Aβ which i) safely preserves Aβ-mediated effector function, ii) enhances brain exposure, parenchymal distribution, and target engagement, and iii) substantially reduces ARIA-like lesions in a mouse model with parenchymal and vascular amyloid pathology. We propose that the improved ARIA profile with ATV^cisLALA^:Aβ is a consequence of its distinct route of entry and biodistribution in the brain compared to standard anti-Aβ.

## Results

### Asymmetrical cisLALA Fc mutations mitigate TfR-mediated, but maintain Fab-mediated, ADCC and CDC *in vitro*

TfR-mediated reticulocyte loss is driven by simultaneous binding to TfR and FcɣR on TfR-expressing early erythroid progenitors (Couch et al., 2013). We previously introduced effector-reducing mutations in the Fc of ATV (L234A/L235A or ‘LALA’) to significantly reduce binding to FcɣRs (Kariolis et al., 2020). As ATV molecules are generated using knob-in hole technology (Merchant et al., 1998), it provided an opportunity to determine whether we can safely introduce FcɣR binding on only one side of the Fc while maintaining TfR binding. To accomplish this, we engineered the ATV with LALA mutations on either the knob (ATV^cisLALA^) or hole side (ATV^transLALA^) of the Fc and compared the properties of these variants to a molecule that has symmetrical LALA (ATV^LALA^) as well as an ATV with a wild-type Fc (ATV^WT^) (**Fig**. **1a**).

**Figure 1.**
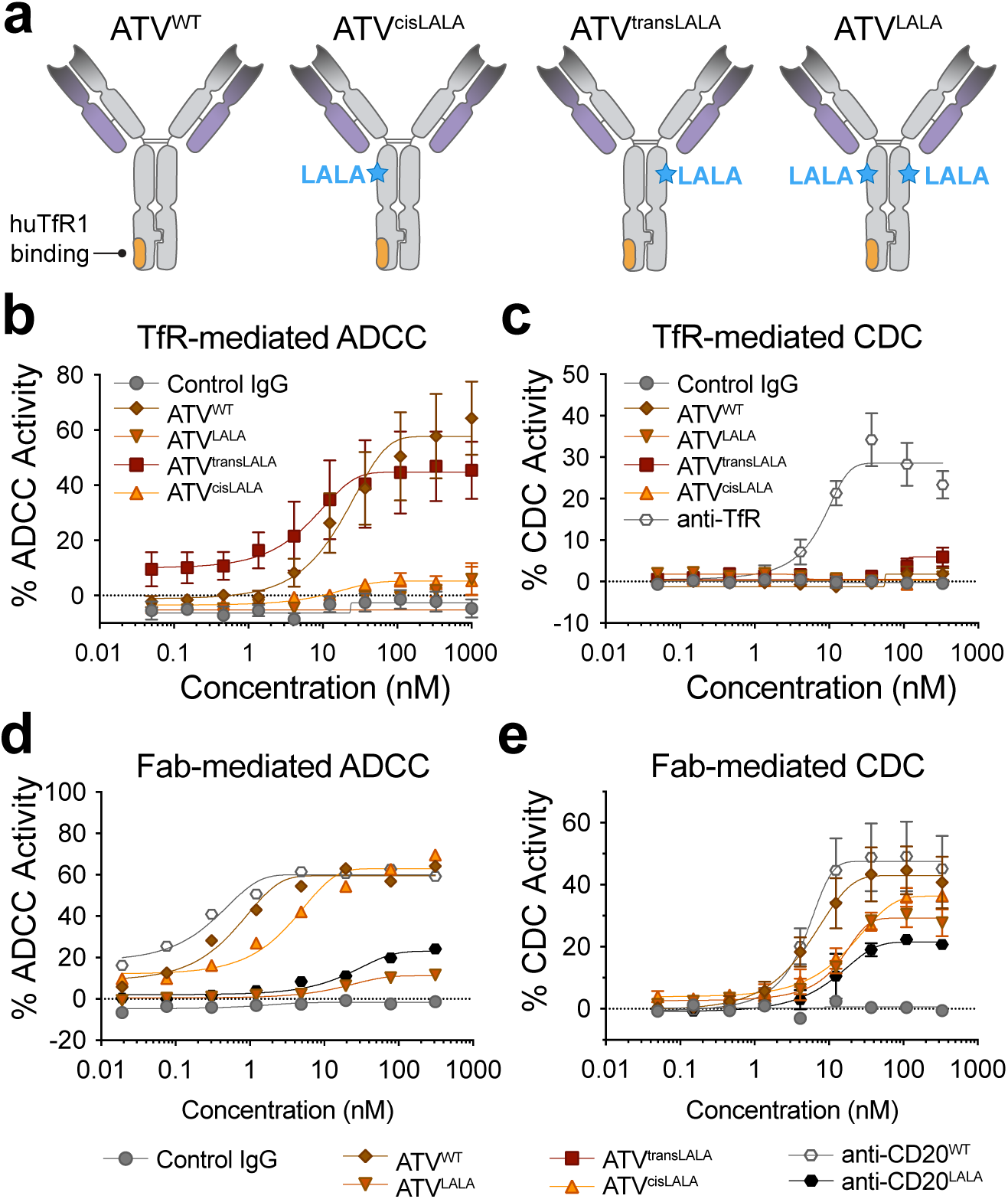
Asymmetrical cisLALA Fc mutations mitigate TfR-mediated but maintain Fab-mediated ADCC and CDC *in vitro*. **a**, Schematics of ATV molecular architectures indicating L234A/L235A (LALA) mutations, if present. **b-c**, In human TfR-expressing cells, ATVs with control BACE1 Fabs were used to evaluate ADCC-induced killing of Ramos cells by isolated human NK cells (**b**) or CDC-induced killing of huTfR-overexpressing CHO cells (**c**). **d**, ATVs with murine (**d**) or human (**e**) CD20 Fabs were used to evaluate Fab-mediated ADCC using mouse A20 lymphoma cells and isolated human NK cells or Fab-mediated CDC using Raji cells. All graphs represent *n*=3 replicate experiments except anti-CD20^LALA^ in (**e**) which had *n*=2 replicate experiments. The ATV affinity for the huTfR apical domain in these experiments was ∼100 nM.

The TfR-mediated antibody-dependent cell cytotoxicity (ADCC) and complement-dependent cytotoxicity (CDC) of these effector function variants were evaluated using ATVs containing the well-characterized BACE1 Fabs previously described (Kariolis et al., 2020). BACE1 Fabs have limited binding to the cells used in this assay, thus isolating the impact of TfR binding. ATV^cisLALA^:BACE1 exhibited a similar attenuation of TfR-mediated ADCC as observed with ATV^LALA^:BACE1, whereas ATV^transLALA^:BACE1 showed comparable level of ADCC as ATV^WT^:BACE1 (**Fig. 1b**). This indicated the ATV^transLALA^ architecture preserved TfR-mediated effector function *in vitro*, possibly due to a maintenance of its orientation on the cell surface for successful FcɣR recruitment, while TfR-bound ATV^cisLALA^ architecture prevented FcɣR binding and TfR-mediated ADCC. Interestingly, none of the ATV molecules were able to elicit CDC on TfR target cells which was clearly present for an anti-TfR mAb (**Fig. 1c**), suggesting that the Fc-binding orientation to TfR likely prevents hexameric C1 interactions that are required to elicit CDC (Diebolder et al., 2014).

We next investigated whether desirable Fab target-mediated effector function of the ATV^cisLALA^ architecture was maintained. To test this, we used murine CD20 Fabs for proof-of-concept, as CD20 antibodies are known to mediate ADCC and CDC on lymphoma cells (Reff et al., 1994). ATV^cisLALA^:CD20 elicited strong ADCC activity on A20 murine lymphoma cells and achieved an E_max_ that was comparable to both ATV^WT^:CD20 and an anti-CD20 mAb with WT huIgG Fc, though it exhibited a slight shift in EC_50_ (**Fig. 1d**). As expected, the effectorless ATV^LALA^:CD20 and anti-CD20 with LALA mutations had significantly blunted ADCC activity. All molecules elicited CDC activity using human CD20 Fabs, with slight attenuation observed for ATV^cisLALA^:CD20, ATV^LALA^:CD20 and anti-CD20^LALA^ mAb (**Fig. 1e**). Taken together, these *in vitro* effector function assays suggested that ATV^cisLALA^ has the potential to eliminate TfR-mediated effector function for hematological safety, while preserving Fab-mediated effector function.

### ATV^cisLALA^ architecture mitigates *in vivo* hematology liability in mice

To determine whether mitigation of *in vitro* TfR-mediated ADCC with ATV^cisLALA^ translates *in vivo*, we formatted ATV effector variants onto anti-Aβ Fabs and evaluated the hematology impacts after systemic dosing in mice. Strikingly, ATV^WT^:Aβ resulted in a near complete depletion of circulating reticulocytes 24 hours after a single 10 mg/kg IV dose in TfR^mu/hu^ KI mice, while ATV^cisLALA^:Aβ and ATV^LALA^:Aβ had no significant reduction in reticulocytes in the same paradigm (**Fig. 2a**). A similar differential effect on reticulocytes was observed in the bone marrow of these mice, where ATV^cisLALA^ and ATV^LALA^ did not have an impact on reticulocytes compared to the anti-Aβ control, while ATV^WT^:Aβ showed a significant reduction (**Fig. 2b**).

**Figure 2.**
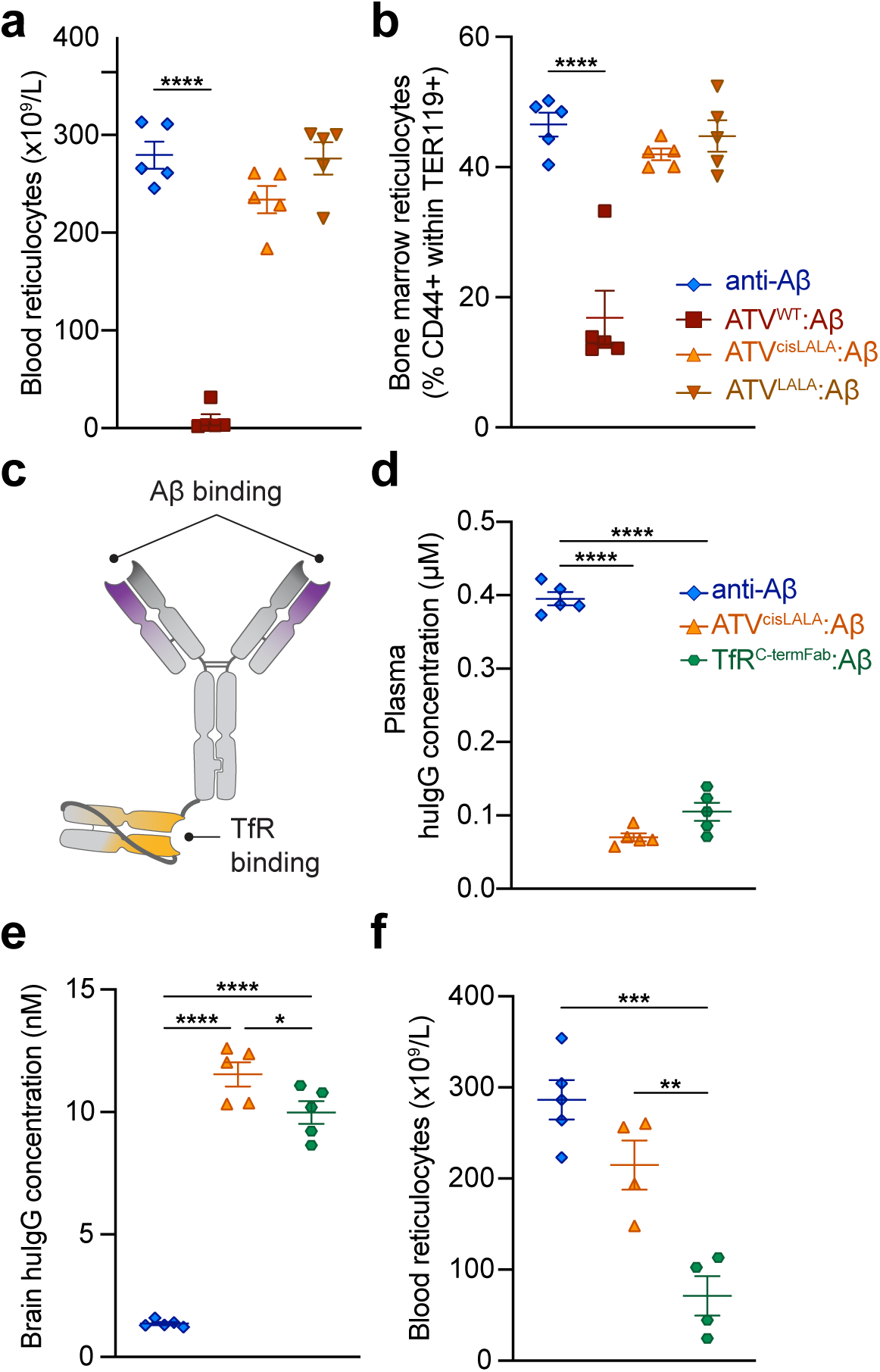
ATV^cisLALA^, but not TfR^C-term^ ^Fab^:Aβ, mitigates *in vivo* reticulocyte loss in mice. **a-b**, Circulating blood (**a**) or bone marrow (**b**) reticulocytes in TfR^mu/hu^ KI mice 24 h after a single 10 mg/kg IV dose of anti-Aβ or ATV:Aβ variants. **c**, An alternative architecture TfR^C-term^ ^Fab^:Aβ generated for comparison of effector function-mediated reticulocyte response. **d-f**, Plasma (**d**) and brain (**e**) concentrations as measured by ELISA, and blood reticulocytes (**f**) 24 h after a single 10 mg/kg IV dose of anti-Aβ, ATV^cisLALA^:Aβ, or TfR^C-term^ ^Fab^:Aβ in TfR^mu/hu^ KI mice. Graphs display mean ± SEM. Only significant differences indicated on graphs; (**a,b,d-f**) one-way ANOVA with multiple comparisons; (**a-b**) Dunnett’s post hoc versus anti-Aβ control, \*\*\**p* < 0.001; (**d-f**) Tukey’s post hoc comparing all groups, \**p* < 0.05, \*\**p* < 0.01, \*\*\**p* < 0.001, \*\*\*\**p* < 0.0001. The ATV and TfR^C-term^ ^Fab^ affinity in these experiments was ∼500 nM (**a-b**, huTfR apical domain) or ∼40-50 nM (**d-f**, full-length huTfR).

Other TfR-based CNS antibody delivery platforms have explored alternative architectures to increase brain exposure while maintaining FcɣR binding (Niewoehner et al., 2014; Weber et al., 2018). It has previously been suggested that appending a C-terminal TfR binding domain onto wild-type huIgG Fc may sterically hinder FcɣR binding and prevent immune cell engagement while bound to TfR, thus preventing TfR-mediated cell killing but retaining Fab-mediated effector function (Weber et al., 2018). To evaluate how this architecture compares to ATV^cisLALA^, we generated an anti-Aβ antibody with full effector function with a C-terminal TfR-binding Fab (**Fig. 2c**) and compared it to the same Aβ antibody with an ATV^cisLALA^ architecture and similar huTfR binding affinity. TfR^C-term^ ^Fab^:Aβ and ATV^cisLALA^:Aβ exhibited comparable target-mediated clearance from plasma (**Fig. 2d**) and both showed a similarly significant enhancement in brain exposure in whole brain 24 hours after a single 10 mg/kg IV dose (**Fig. 2e**), suggesting the differences between the two molecule architectures did not differentially impact overall *in vivo* exposure. However, while ATV^cisLALA^:Aβ had no significant impact on circulating reticulocytes, TfR^C-term^ ^Fab^:Aβ treated mice experienced a significant reduction in circulating reticulocytes (**Fig. 2f**), consistent with other recent reports using a similar TfR-binding architecture with full effector function (Edavettal et al., 2022). These results suggest that steric hindrance was insufficient to protect against effector-function driven loss of TfR-expressing reticulocytes and that the ATV^cisLALA^ architecture provides a unique opportunity to mitigate TfR-mediated, effector-function driven hematology impact in mice while retaining TfR engagement.

### ATV^cisLALA^:Aβ retains ability to induce microglia phagocytosis *ex vivo* and reduce plaques *in vivo*

We next investigated whether the cisLALA mutations retain the ability to induce anti-Aβ-mediated phagocytosis, as it is a critical component of amyloid plaque removal (Bamberger et al., 2003; Bard et al., 2000). Phagocytosis was evaluated using flow cytometry by incubating freshly dissociated TfR^mu/hu^ KI mouse brain cells with a preparation containing a mix of fluorescein (FAM)-labeled Aβ aggregates (**Supplemental Fig. 1a**), along with either control IgG, anti-Aβ (WT Fc), ATV^WT^:Aβ, ATV^cisLALA^:Aβ, or ATV^LALA^:Aβ. Cells were then assessed by flow cytometry to detect the number of FAM-Aβ-positive and FAM-Aβ-negative microglia (**Supplemental Fig. 1d**). This *ex vivo* paradigm was designed to specifically measure the microglial phagocytosis of these molecules and avoids confounding effects of differential brain exposure and biodistribution between non-ATV and ATV molecules. Analysis revealed comparable significant increases in the percentage of Aβ-positive microglia for anti-Aβ, ATV^cisLALA^:Aβ, and ATV^WT^:Aβ treated groups compared to control IgG (**Fig. 3a-b**). In contrast, ATV^LALA^ treated cells exhibited a significantly lower level of phagocytosis compared to other groups, consistent with a reduction in functional FcɣR engagement. A lack of FAM-Aβ signal in microglia at 4°C (**Supplementary Fig.1b-c**) further confirmed the observed increase in fluorescence intensity was due to phagocytic uptake, rather than simply binding to the cell surface.

**Figure 3.**
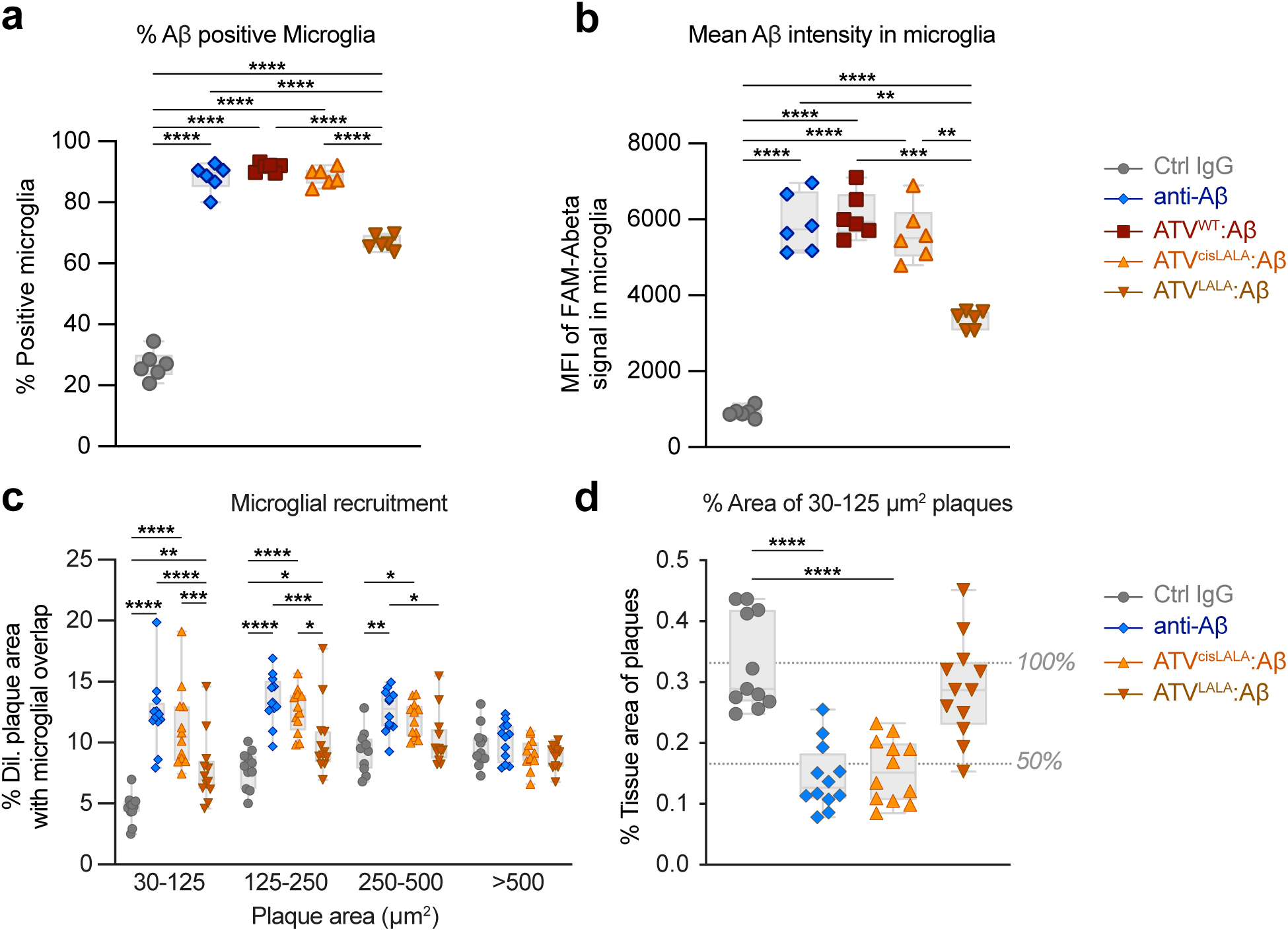
ATV^cisLALA^:Aβ retains the ability to induce microglia phagocytosis *ex vivo* and recruits microglia to clear plaques *in vivo.* **a-b**, Freshly perfused brains from TfR^mu/hu^ KI mice were dissociated and brain cells were incubated with FAM-labeled Aβ and either control IgG, anti-Aβ, or ATV:Aβ variants at 37°C. Uptake of labeled Aβ into live microglia was measured using flow cytometry to assess *ex vivo* phagocytosis. The percentage of live microglia with positive Aβ signal (**a**) and the mean fluorescence intensity of Aβ signal in live microglia (**b**) were compared. **c-d**, *In vivo* phagocytosis was assessed using an acute multidose paradigm (50 mg/kg Q3Dx4, IP) in mixed sex 5xFAD; TfR^mu/hu^ KI mice to achieve high brain exposure for all groups by the terminal time point of 12 days. CD68-positive microglial recruitment to plaques (**c**), measured by overlap with plaques and a small, dilated region immediately surrounding plaques, and percent tissue area of small Aβ plaques (**d**). Only significant differences indicated on graphs; (**a,b,d**) one-way ANOVA or (**c**) two-way ANOVA with multiple comparisons; (**a-c**) Tukey’s post hoc comparing all groups, \**p* < 0.05, \*\**p* < 0.01, \*\*\**p* < 0.001, \*\*\*\**p* < 0.0001 (**d**) Dunnett’s post hoc versus Ctrl IgG, \*\*\*\**p* < 0.0001. The huTfR apical domain affinity in these experiments is ∼1200 nM (**a-b**) or ∼500 nM (**c-d**).

We next crossed the 5xFAD transgenic AD mouse model (Oakley et al., 2006) with TfR^mu/hu^ KI mice (Kariolis et al., 2020) for functional *in vivo* evaluation of ATV:Aβ variants. To assess whether the ATV^cisLALA^ architecture retains full ability to recruit microglia to Aβ plaques, we used an acute multidose paradigm to achieve similar total brain exposures between the ATV^LALA^:Aβ, ATV^cisLALA^:Aβ, and anti-Aβ groups (**Supplementary Fig. 1e**) to focus on the role of the Fc mutations on microglial recruitment. Interestingly, although bulk brain lysate levels were similar across all groups, the two ATV molecules had greater immunodecoration of plaques compared to anti-Aβ indicating improved target engagement (**Supplementary Fig. 1f-g**).

Aβ plaques were binned by size to elucidate differences in microglial recruitment that might arise between newer or more established plaques. Colocalization of the activated microglial marker CD68 with Aβ plaques and their immediate surroundings revealed similar microglial recruitment in mice treated with ATV^cisLALA^:Aβ compared to anti-Aβ with WT Fc, with a notable deficit for the effectorless ATV^LALA^:Aβ (**Fig. 3c**, **Supplementary Fig. 1h**). This effect was the most significant for smaller plaque sizes and maintained a similar trend for larger plaques. Surprisingly, despite the relatively acute treatment duration, we also observed that both ATV^cisLALA^:Aβ and anti-Aβ showed a similar and significant reduction in burden of the smallest plaques (30-125 mm^2^) compared to control, whereas ATV^LALA^:Aβ treatment resulted in no significant plaque reduction (**Fig. 3d**). Together, these data suggest that preservation of effector function is critical to recruit microglia to plaques and contributes to the clearance and/or prevention of small plaques in this short time span. Importantly, ATV^cisLALA^:Aβ preserves sufficient effector function to enable this process.

### ATV^cisLALA^ has superior brain uptake and target engagement

There have been limited studies evaluating brain uptake of the ATV platform in mouse models with plaque pathology (van Lengerich et al., 2023). We therefore sought to determine whether plaque burden has an impact on brain uptake of ATV^cisLALA^:Aβ at therapeutically relevant dose levels in 4-month-old 5xFAD; TfR^mu/hu^ KI mice. NP40-soluble brain lysate concentrations of ATV^cisLALA^:Aβ were approximately 5-8 times higher for ATV^cisLALA^:Aβ compared to anti-Aβ at all dose levels 24 hours post-dose (**Fig. 4a**). Moreover, the percent brain:plasma was significantly higher for ATV^cisLALA^:Aβ, indicating a more efficient transport into the brain compared to anti-Aβ in both models (**Fig. 4b**; **Supplementary Fig. 2a**). Immunodetection of the administered anti-Aβ or ATV^cisLALA^:Aβ huIgG in brain sections also revealed significantly greater target engagement of ATV^cisLALA^:Aβ with Aβ plaques (**Fig. 4c-e**). CSF concentrations of anti-Aβ and ATV^cisLALA^:Aβ were similar and exhibited dose-dependence (**Supplementary Fig. 2b**) despite significant differences in brain lysate concentration, a finding consistent with previous reports suggesting CSF generally underestimates brain concentrations of TfR-binding ATV molecules (Kariolis et al., 2020) and consistent with predominant transport of ATVs across the BBB (Khoury et al., 2024).

**Figure 4.**
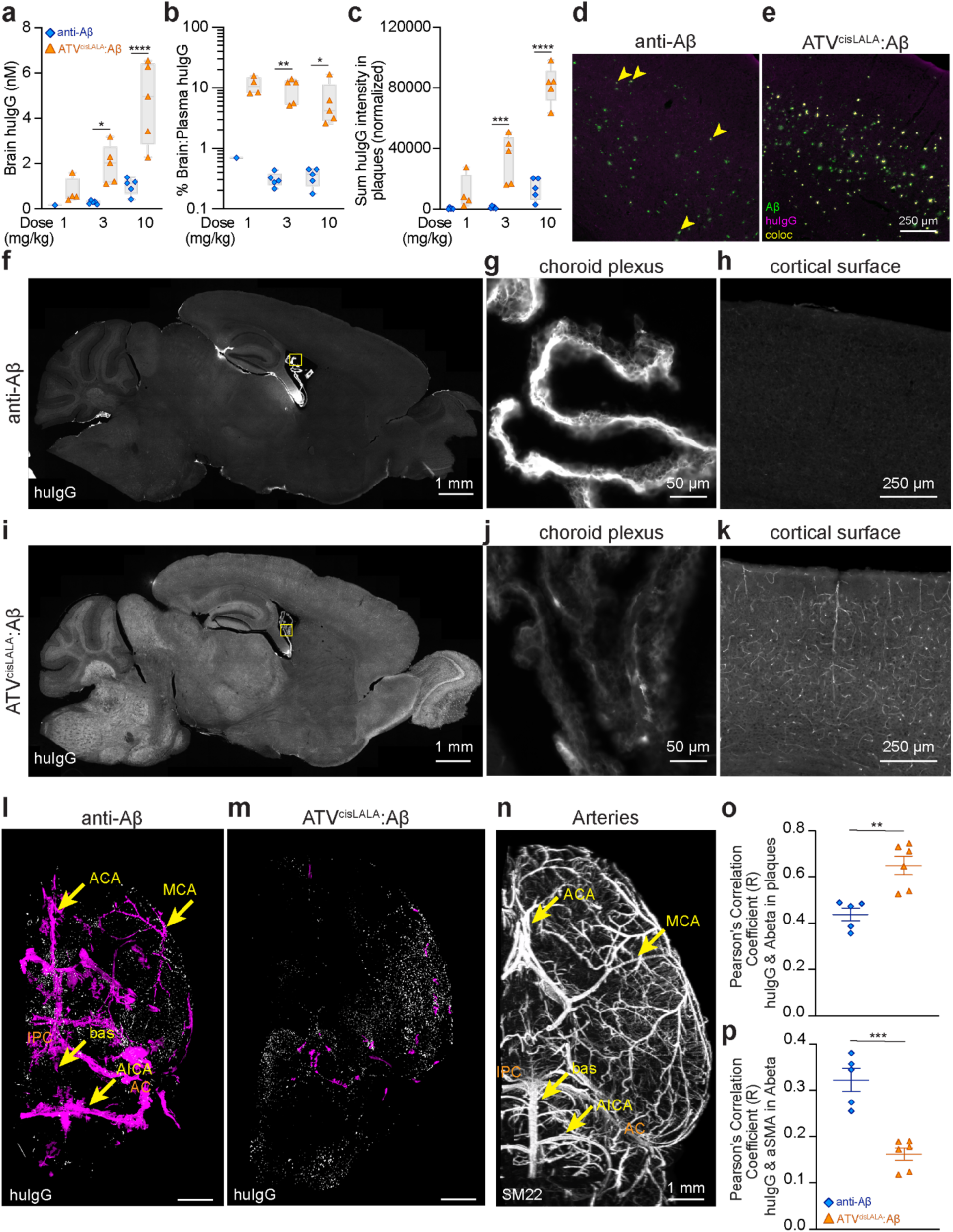
ATV^cisLALA^ has superior brain uptake, biodistribution, and target engagement and lesser association with arteries in mice. **a-e**, 5xFAD; TfR^mu/hu^ KI mice received a single IV dose of 1, 3, or 10 mg/kg anti-Aβ or ATV^cisLALA^:Aβ. huIgG concentrations were measured by ELISA. Brain lysate concentration 24 h post-dose (**a**) was used to calculate the percentage of brain concentration to plasma concentration (**b**). In (**a-b**) *n*=4/5 samples below the limit of quantification for anti-Aβ. Quantification of plaque decoration by huIgG (**c**) and representative widefield images from the cortex following 10 mg/kg anti-Aβ (**d**) or ATV^cisLALA^:Aβ (**e**). **f-k**, Representative widefield images of huIgG distribution 24 h after anti-Aβ (**f-h)** or ATV^cisLALA^:Aβ (**i-k**) after a single 25 mg/kg IV dose in TfR^mu/hu^ KI mice. Magnified examples of huIgG signal in the choroid plexus and cortex from anti-Aβ group (**g-h**) or ATV^cisLALA^:Aβ group (**j-k**). **l-n**, 5xFAD; TfR^mu/hu^ KI mice were IV dosed with 10 mg/kg of AF750-conjugated ATV^cisLALA^:Aβ or anti-Aβ and perfusion-fixed 24 h post-dose. Due to low signal-to-noise with direct detection of the AF750 fluorophore label, whole hemibrains were also immunostained for huIgG signal. One hemisphere evaluated per staining set; *n*=2/group in **(l)** and (**m**) (no additional markers), *n*=6/group in (**o**) and (**p**) (additional immunodetection of Aβ and αSMA, **Supplementary Fig. 3**). huIgG signal was segmented based on size (<50 μm^3^ white, >50 μm^3^ magenta) for anti-Aβ (**l**) and ATV^cisLALA^:Aβ (**m**). In a separate experiment to illustrate anatomical pattern of surface arteries, fixed mouse brains were stained with SM22 to detect smooth muscle cells of arteries (**n**). **o-p**, Aβ-positive objects (plaques and vascular amyloid) were segmented based on Aβ immunodetection and the Pearson’s Correlation between huIgG and Aβ signal within Aβ-positive objects (**o**) or huIgG and alpha-smooth muscle actin (αSMA) signal in Aβ-positive objects (**p**). Anterior cerebral artery (ACA), middle cerebral artery (MCA), basilar artery (bas), anterior inferior cerebellar artery (AICA), interpeduncular cistern (IPC), ambient cistern (AC). Only significant differences indicated on graphs; (**a-c**) one-way ANOVA; Sidak’s multiple comparisons test paired by dose level, \**p* < 0.05, \*\**p* < 0.01, \*\*\**p* < 0.001, \*\*\*\**p* < 0.0001; (**o-p**) two-tailed t-test on Fisher transformed correlation coefficients, \*\**p* < 0.01, \*\*\**p* < 0.001. The ATV affinity for huTfR apical domain in these experiments is ∼500 nM (**a-e**, **l-p**) or ∼1200 nM (**i-k**).

We hypothesized that the TfR-mediated entry across the BBB would additionally result in highly differentiated distribution of ATV^cisLALA^:Aβ in the brain compared to standard antibodies. Spatial distribution analysis of anti-Aβ and ATV^cisLALA^:Aβ in TfR^mu/hu^ KI mice one day after a single dose revealed that anti-Aβ signal was predominantly associated with cells and brain regions bordering the CSF compartment, including the choroid plexus, periventricular brain regions, pial brain surfaces, leptomeningeal cells, and perivascular components, particularly along the ventral brain and CSF cisterns (**Fig. 4f-h**). In sharp contrast, ATV^cisLALA^:Aβ signal was distributed broadly throughout the entire sagittal brain section, with signal present in the choroid plexus (appearing as punctate intracellular signal) and prominently distributed across brain regions associated with the brain vasculature and parenchyma (**Fig. 4i-k**). Thus, ATV^cisLALA^:Aβ not only increased overall brain concentrations but also resulted in a broader, whole-brain distribution that included deep brain regions, consistent with our findings for non-targeted antibodies compared to ATVs (Khoury et al., 2024).

Given the striking localization of anti-Aβ with CSF-associated spaces, we further compared the distribution of ATV^cisLALA^:Aβ and anti-Aβ in a plaque-bearing mouse model using tissue clearing and 3D imaging. This approach enabled us to better visualize brain surface structures (i.e., leptomeningeal arteries and perivascular spaces) where CAA, a common vascular pathology in AD, is predominantly localized (Thal et al., 2008). Engagement of CAA by anti-Aβ antibodies is hypothesized to be a trigger for the perivascular inflammatory response which likely underlies ARIA, adverse events observed in patients treated with amyloid immunotherapy (Doran & Sawyer, 2024; Greenberg et al., 2020; Keable et al., 2016; Sveikata et al., 2022; Weller et al., 1998). We performed 3D light sheet imaging on iDISCO-cleared whole hemibrains from 3-3.5 month old 5xFAD; TfR^mu/hu^ KI mice dosed with anti-Aβ or ATV^cisLALA^:Aβ. One hemibrain was immunostained for huIgG and the signal was segmented by object size (smaller or larger than 50 μm^3^). Anti-Aβ signal (**Fig. 4l**, **Supplementary Fig. 3a**) showed substantial localization to the larger, connected volumes on the brain surface in a pattern consistent with major surface arteries (Fig 4n), presumably associated with arterial perivascular spaces and leptomeningeal cells. Anti-Aβ signal was also prominently localized adjacent to CSF cisterns; overall there was relatively little association with small puncta (presumed Aβ plaques, **Supplementary Fig. 3a**). In contrast, ATV^cisLALA^:Aβ (**Fig 4m**) was predominantly associated with object sizes smaller than 50 μm^3^ (putative parenchymal plaques) and showed less association with leptomeningeal arteries (**Supplementary Fig. 3b**). The other hemibrain was immunostained for Aβ to confirm the identify of plaques, alpha smooth muscle actin (αSMA) to identify vascular smooth muscle cells predominantly associated with arteries, and huIgG for the dosed molecules (**Supplementary Fig. 3**). Despite the relatively young age of the 5xFAD; TfR^mu/hu^ KI mice, 3D imaging revealed apparent CAA (i.e., Aβ present around smooth muscle cells, **Supplementary Fig. 3c-f**) associated with large surface arteries in most animals. Quantification of huIgG signal within plaques revealed a significantly greater correlation of huIgG and Aβ in plaques for ATV^cisLALA^:Aβ compared to anti-Aβ (Fig 4o), suggesting greater target engagement by ATV. In contrast, ATV^cisLALA^:Aβ treated mice had significantly less correlation of huIgG and αSMA within Aβ positive objects (i.e., vascular amyloid) compared to anti-Aβ (Fig 4p, **Supplementary Fig. 3c-f**), suggesting anti-Aβ is more commonly associated with arterial amyloid (i.e., CAA).

### ATV^cisLALA^:Aβ significantly reduces the incidence of ARIA-like lesions in mice

We hypothesized that the higher vascular and perivascular anti-Aβ signal that co-localized with CAA might well contribute to increased inflammatory responses and a greater risk of ARIA following anti-Aβ treatment compared to ATV^cisLALA^:Aβ. To test this, we employed the chimeric (ch) 3D6 anti-Aβ antibody for additional studies, as it has been well documented to induce CAA-associated microhemorrhages in mouse models of AD (Demattos et al., 2012; Racke et al., 2005; Schroeter et al., 2008; Taylor et al., 2023; Zago et al., 2013), and a robust mouse ARIA model was recently described using this molecule (Bussiere, 2024). 5xFAD; TfR^mu/hu^ KI mice were administered weekly 3 mg/kg of either ^ch^3D6^WT^ or ATV^cisLALA^:3D6 (**Fig. 5a**). An additional ATV^cisLALA^:3D6 group was dosed at 1.75 mg/kg to approximately match the expected brain AUC of ^ch^3D6^WT^. Because no ARIA-like magnetic resonance imaging (MRI) events were detected by week 4 for any of the groups, dose levels were increased at week 5 for all groups by ∼3x (i.e., 1.75 to 5 mg/kg and 3 to 10 mg/kg). Strikingly, the rate of ARIA-like MRI lesions increased in the ^ch^3D6^WT^-treated group, with all mice (10 out of 10) displaying at least one ARIA-like event by week 8 (**Fig. 5b**). In contrast, only one mouse in the dose-matched ATV^cisLALA^:3D6 displayed an MRI lesion at week 9 (which persisted until the end of the study at week 12), despite the expected higher brain exposure in this group compared to the ^ch^3D6^WT^ group. Importantly, no ARIA-like events occurred in the brain exposure-matched ATV^cisLALA^:3D6 group.

**Figure 5.**
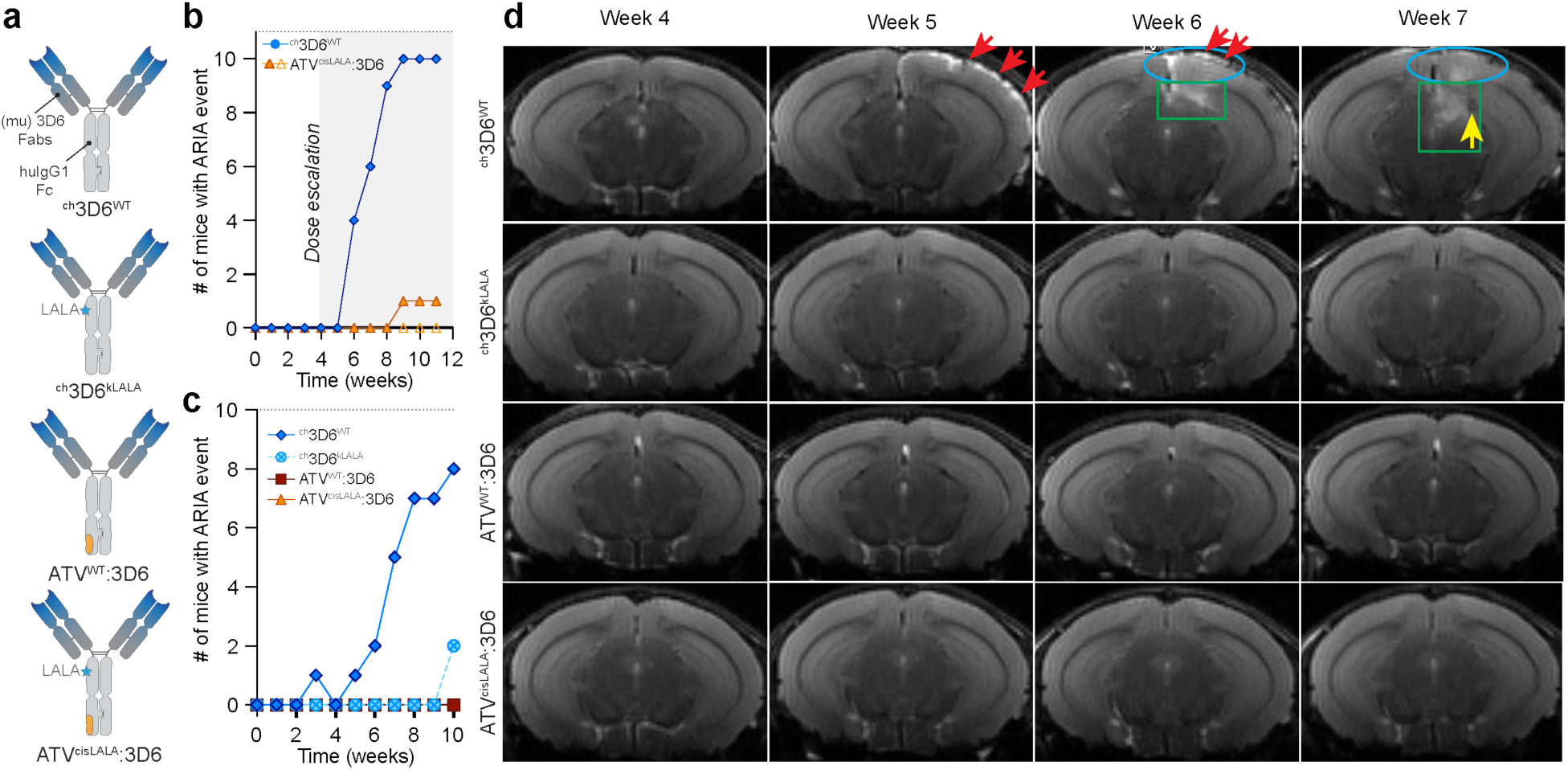
TfR-mediated brain delivery reduces incidence of ARIA-like lesions in 5xFAD; TfR^mu/hu^ KI mice. **a**, Schematics of the molecules designed to assess roles of effector function and TfR-mediated brain uptake in occurrence of ARIA. **b**, Incidence of ARIA-like lesions determined by weekly T2w MRI scans in male 5xFAD; TfR^mu/hu^ KI mice treated with ^ch^3D6^WT^ and dose-matched (closed triangle) or exposure-matched (open triangle) ATV^cisLALA^:3D6 starting at 3 or 1.75 mg/kg and increasing to 5 or 10 mg/kg at the 5^th^ dose (*n*= 11 mice/group). **c**, Incidence of ARIA-like lesion determined by weekly T2w MRI scans in mixed sex 5xFAD; TfR^mu/hu^ KI mice treated with ^ch^3D6 (10 mg/kg) or exposure-matched ATV variants (5 mg/kg) (*n*= 10 mice/group). **d**, Representative scans from study in (**c**) collected using a T2w MRI sequence. Several types of ARIA-like lesions were detected in mice treated with ^ch^3D6^WT^: hyperintense convexity lesion (red arrow, week 5-6), hyperintense diffuse cortical (blue oval, week 6-7), sub-cortical lesion (green rectangle, week 6-7), and hypointense focal sub-cortical lesion (yellow arrow, week 7).

We next generated two additional molecules for a second study to better separate the contribution of ATV-mediated delivery versus effector function in the mitigation of ARIA events: ^ch^3D6^kLALA^, with asymmetric LALA mutations on one strand of the huIgG Fc to mirror ATV^cisLALA^, and ATV^WT^:3D6, which had full effector function owing to its wild-type huIgG Fc (**Fig. 5a**). To rule out a potential contribution of differential brain exposure to the results, dose levels were selected to approximately achieve similar brain exposure over the duration of this second study for each of four groups. 5xFAD; TfR^mu/hu^ KI mice treated weekly with 10 mg/kg ^ch^3D6^WT^ developed ARIA-like lesions as early as week 3, with 8 out of 10 mice displaying at least one lesion by the end of the study (week 10; **Fig. 5c**). The MRI score, accounting for size and severity of lesions, for ^ch^3D6^WT^-treated animals was greater than any of the other treatment groups (**Supplemental Table 1**). ^ch^3D6^kLALA^ treatment resulted in 2 out of 10 mice developing ARIA-like lesions in the last week of the study. Strikingly, none of the mice treated weekly with 5 mg/kg ATV^WT^:3D6 or ATV^cisLALA^:3D6 developed any ARIA-like lesions throughout the duration of the study (**Fig. 5c**, **Supplemental Table 1**). ARIA-like lesions were associated with both hyperintensities and hypointensities in scans collected using a T2w MRI sequence (Fig 5d). T2-hyperintense convexity lesions or T2-hyperintense cortical lesions were located within the meningeal space or in the depth of the cortex, respectively. T2-hyperintense focal cortical lesions were visualized as bright and well-circumscribed signals, whereas T2-hyperintense diffuse lesions were visualized as attenuated signal and were observed to occupy a large volume on T2w scans. T2-hypointense lesions were usually small in size, and well-circumscribed within the brain parenchyma. They were less frequent than T2-hypertense lesions and occurred as individual lesions or as an evolution of T2-hyperintense lesions. Taken together, these MRI findings demonstrate that ATV^WT^ and ATV^cisLALA^ architectures effectively mitigated ARIA-like events.

### Histopathology shows the ATV format is associated with lower vascular inflammation and disruption

Blinded histopathologic examination of coronal brain sections from ^ch^3D6^WT^-treated mice in the expanded study (**Fig. 5c**) revealed frequent leptomeningeal foci of Grade 3-4 vascular inflammation, perivascular and diffuse-parenchymal albumin immunoreactivity and perivascular hemosiderin, in contrast to the complete absence of these pathologies in untreated mice (**Fig. 6a-d**). ^ch^3D6^kLALA^-treated mice and ATV^WT^:3D6-treated mice showed fewer foci of mild-to-moderate meningovascular inflammation with only sparse foci of Grade 4 vascular inflammation and vascular mural compromise. Importantly, ATV^cisLALA^:3D6-treated mice had a near absence of meningovascular inflammation; only rare foci of Grade 1 meningovascular inflammation were observed and there was an absence of abnormal vascular permeation and perivascular hemosiderin (**Fig. 6a-d**). Though far less prevalent than foci of meningovascular inflammation, cortical microinfarcts were present in most ^ch^3D6^WT^-treated mice and trended lower in ^ch^3D6^kLALA^-treated mice (**Fig. 6e**). Notably, ATV^WT^:3D6-treated mice showed no microinfarcts in the sampled histologic sections. Only one miniscule remote microinfarct was observed in one of the ATV^cisLALA^:3D6-treated mice, and its histologic features (i.e., astrogliosis with lack of macrophage activity) suggested it may have preceded ATV^cisLALA^:3D6 treatment (**Fig. 6e**, ‡). A heatmap of these pathologic changes reveal concordance of these vascular pathologies in individual mice among the ^ch^3D6^WT^-treated and ^ch^3D6^kLALA^-treated groups, and absent from the ATV and untreated groups (**Fig. 6f**). Similar reductions in meningovascular inflammation, vascular mural leakage, and microinfarct pathologies were observed in the initial study (**Fig. 5b**) comparing ^ch^3D6 with equal or exposure-matched doses of ATV^cisLALA^:^ch^3D6 (**Supplementary Fig. 4**). Taken together, these MRI and histopathology data suggest that ATV robustly reduces the incidence of ARIA-like lesions and underlying vascular inflammation in a mouse model of AD.

**Figure 6.**
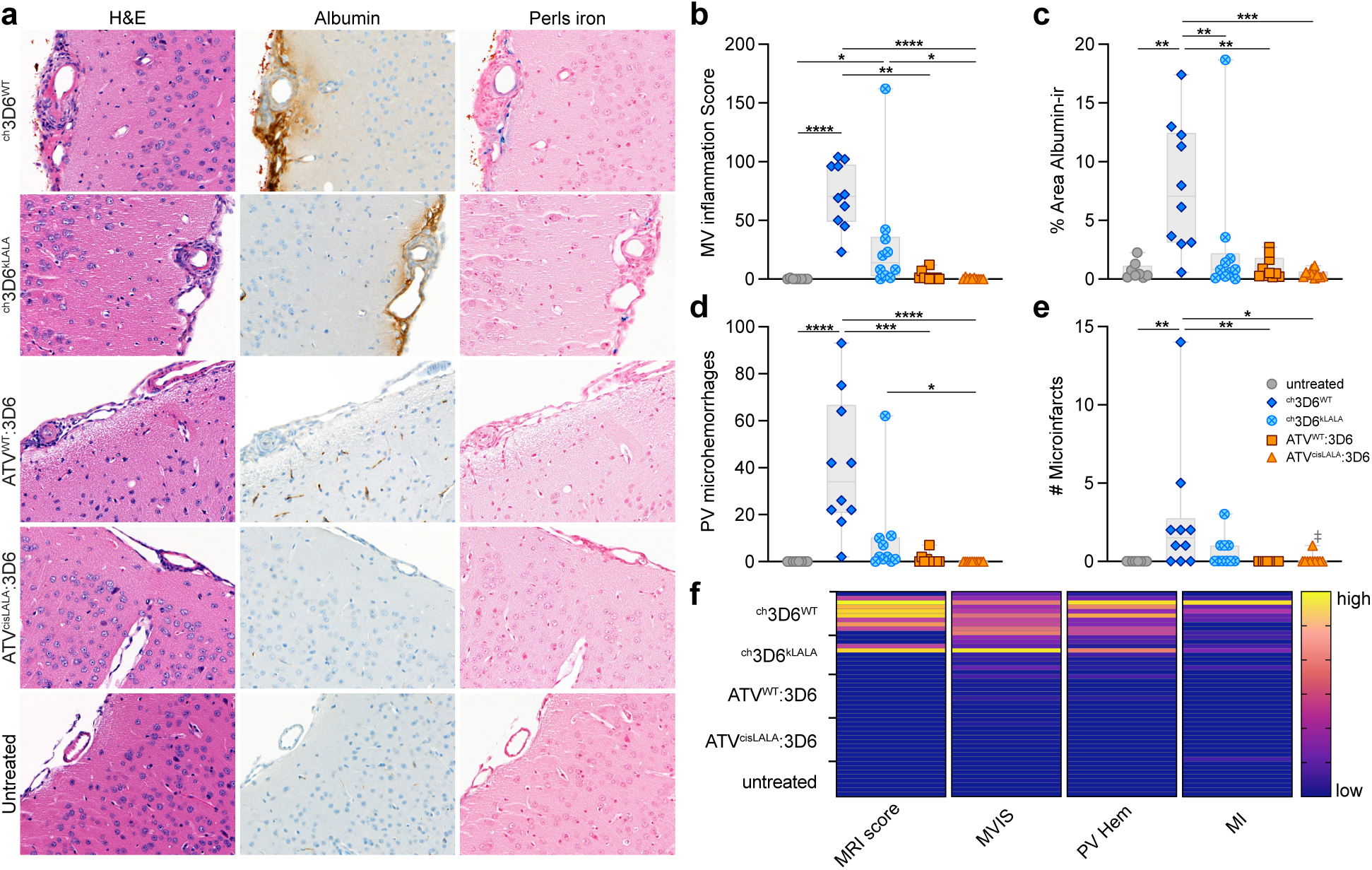
^ch^3D6-induced meningovascular pathology is mitigated by both TfR-mediated brain uptake and asymmetrical FcɣR binding. **a-f**, Blinded histopathological assessment of brains from mixed sex 5xFAD; TfR^mu/hu^ KI mice treated with ^ch^3D6 (10 mg/kg) or exposure-matched ATV variants (5 mg/kg). Representative images of H&E, albumin immunoreactivity, and Perls iron from the from study in **Fig. 5c**. **b-e,** Plots of summated meningovascular inflammation scores (**b**), cortical albumin immunoreactivity (**c**), Perls-reactive perivascular hemosiderin (**d**) and microinfarcts (**e**) demonstrating reductions in these ^ch^3D6^WT^-induced pathologies with ATV and kLALA or cisLALA modifications. ‡The one microinfarct identified in the blinded review of the ATV^cisLALA^:3D6 group demonstrated histologic features of remote occurrence (astrogliosis without macrophage activity) and lacked associated meningovascular inflammation that suggest it was not induced by the treatment. **f,** Heatmap demonstrating the animal-by-animal correlations of ARIA-like MRI lesions and histopathologic changes; heatmap scales in order left to right: 0-10, 0-170, 0-100, 0-15. Only significant differences indicated on graphs; (**c**) one-way ANOVA with multiple comparisons or (**b, d, e**) Kruskal-Wallis test used when D’Agostino & Pearson normality test failed for at least one group; (**c**) Tukey’s post hoc comparing all groups, \*\**p* < 0.01, \*\*\**p* < 0.001, (**b, d, e**) Dunn’s post hoc comparing all groups \**p* < 0.05, \*\**p* < 0.01, \*\*\**p* < 0.001, \*\*\*\**p* < 0.0001. The ATV affinity for huTfR apical domain in these experiments is ∼1200 nM.

## Discussion

Here, we demonstrate that the TfR-targeted ATV^cisLALA^ platform improves brain exposure, target engagement, and parenchymal brain biodistribution compared to an unmodified anti-Aβ antibody in a mouse model of amyloid deposition. The unique asymmetrical huIgG1 Fc engineering mitigates TfR-mediated hematology liabilities while preserving the ability to engage microglia and enable phagocytosis of Aβ. Our results also reveal an important additional benefit of TfR-enabled delivery, namely differentiated biodistribution of ATV^cisLALA^:Aβ away from large caliber arteries and arterioles where anti-Aβ tends to be concentrated. The difference in their peri-arterial biodistribution (putative sites of CAA) likely contributes to the almost complete mitigation of microhemorrhages, meningovascular inflammation, and microinfarcts by ATV^cisLALA^:Aβ compared to anti-Aβ in a preclinical mouse model of ARIA. Our results suggest that not only does ATV-mediated delivery significantly improve drug biodistribution and target engagement throughout the brain but, critically, its unique biodistribution may also protect against vascular damage observed in the context of ARIA with amyloid immunotherapies.

Previous studies demonstrated that targeting TfR with wild-type Fc IgGs having full effector function can induce reticulocyte loss (Couch et al., 2013; Edavettal et al., 2022); as a result, effector-mitigating mutations have been traditionally used for TfR-based BBB-crossing platforms (Edavettal et al., 2022; Kariolis et al., 2020; Logan et al., 2021; Ullman et al., 2020; van Lengerich et al., 2023). The ATV^cisLALA^ architecture offers a unique engineering solution to safely enable targets requiring effector function for efficacy, as our results demonstrate that it prevents TfR-mediated, effector function-driven cell killing of reticulocytes cells in mice. ATV^cisLALA^ also appeared to be more protective with respect to hematological effects compared to another effector-enabling TfR-based delivery platform currently being pursued for anti-Aβ (Weber et al., 2018). Importantly, ATV^cisLALA^:Aβ fully preserved Fab-mediated effector function critical to the molecule’s ability to recruit microglia and enable Aβ phagocytosis both *ex vivo* and *in vivo*.

Although recent clinical successes with anti-Aβ mAbs have led to the approval of three therapies for the treatment of AD, most clinically tested anti-Aβ drugs are associated with some level of ARIA (Ostrowitzki et al., 2017; Salloway et al., 2014; Sevigny et al., 2016; R. A. Sperling et al., 2011). While the pathogenic mechanisms underlying the occurrence of ARIA with amyloid therapies are not yet fully understood, it has been hypothesized that ARIA might be related to antibody engagement with CAA and subsequent triggering of an inflammatory response from activated perivascular macrophages, leading to vascular leakage and microhemorrhages (Kozberg et al., 2022; Taylor et al., 2023). ARIA typically occurs early during treatment, is asymptomatic in most cases, and tends to be transient, but some ARIA occurrences may lead to severe adverse events. ARIA complications and monitoring present ongoing challenges to the management of patients taking anti-amyloid antibodies, thus, there is a critical need to identify new ways to reduce ARIA for this class of drugs to reach its full potential. Our experiments reveal that ATV^cisLALA^, a novel TfR-binding format, can dramatically reduce the incidence of ARIA-like adverse events in the 5xFAD mouse model, at least partly owing to its unique biodistribution compared to anti-Aβ, while maintaining the ability to phagocytose and reduce plaque. Additionally, our data suggest that both effector function and TfR-mediated delivery can play important roles in the mitigation of ARIA-like events in this mouse model. Although anti-Aβ^kLALA^ was able to significantly reduce the incidence of ARIA compared to anti-Aβ with WT huIgG, only the ATV^cisLALA^ architecture was able to eliminate MRI and histopathology findings at a brain exposure-matched dose level compared to anti-Aβ. Indeed, even a higher dose of ATV^cisLALA^:Aβ resulted in only a marginal increase in ARIA incidence (**Fig. 5b**). Future preclinical studies to investigate a broader range of dose levels may be warranted to better determine the specific therapeutic window for this approach. Clinical investigation will also be needed to determine whether *APOE ε4* patients, who are at higher risk of ARIA, could significantly benefit from this approach.

Our results are consistent with the model whereby in addition to enhanced brain exposure and target engagement, TfR-mediated brain uptake in capillary beds away from major sites of CAA can be protective against the pathogenic mechanisms underlying ARIA. We utilized multiple imaging methods to reveal that anti-Aβ was largely associated with CSF-communicating compartments within the brain, including the leptomeninges and associated penetrating perivascular compartments, CSF cisterns, and the pial brain surface. This distribution pattern aligns well with evidence that macromolecules such as antibodies circulating in the plasma are typically greatly restricted from transport across the BBB (Abbott et al., 2006; Khoury et al., 2024; St-Amour et al., 2013), with transport across the choroid plexus and/or other extracellular pathways into the CSF likely offering the predominant route of entry (Broadwell & Sofroniew, 1993; Khoury et al., 2024; Rapoport, 1983; Rapoport & Pettigrew, 1979). Unmodified antibodies must then diffuse across leptomeningeal fibroblast layers to reach the parenchymal extracellular spaces of superficial brain regions (Hannocks et al., 2018; Pietilä et al., 2023; Pizzo et al., 2018). Ultimately, further diffusion through the tortuous extracellular spaces to reach deeper sites is greatly restricted by the relative inefficiency of diffusion (Thorne & Nicholson, 2006; Wolak et al., 2015). In contrast, ATV-enabled molecules undertake a markedly different path to brain tissue. Elevated TfR expression on capillaries and postcapillary venules of the mouse brain (Vanlandewijck et al., 2018) coupled with the high density of the neurovascular network (Thorne, 2022) allows for short diffusion distances following transcytosis across brain endothelial cells to enable ATV:Aβ to essentially reach all cells of the brain. Our 3D imaging supports this different route of entry for ATV^cisLALA^:Aβ, which showed predominantly parenchymal distribution, greater plaque target engagement, and far less accumulation around CSF-communicating perivascular spaces, particularly for large surface arteries, consistent with reduced expression of TfR on arterial endothelial cells (Vanlandewijck et al., 2018). Given that accumulation of Aβ around arteries may be up to five-fold higher than around veins in brains of AD patients (Weller et al., 1998), our data revealing less ATV^cisLALA^:Aβ association with arteries (Fig 4) supports a model where the leptomeningeal and perivascular localization of anti-Aβ would be expected to increase the likelihood of anti-Aβ association with CAA, consistent with the high incidence of ARIA-like events observed in our studies. In contrast, a differentiated route of CNS entry for ATV mitigates the incidence ARIA-like events. Indeed, recent interim Phase 1 clinical data with TfR-enabled anti-Aβ trontinemab demonstrated rapid clearance of amyloid plaques with a reduced incidence of ARIA (NCT04639050). These data lend further support to our interpretation from the present study and the clinical utility of a TfR-targeted anti-amyloid approach for AD, although the peripheral safety of using a fully effector positive huIgG will require close monitoring.

Taken together, ATV^cisLALA^ provides a unique and differentiated platform to further improve amyloid immunotherapy efficacy through improved brain exposure, distribution, and target engagement, while mitigating the main adverse events observed with anti-Aβ therapeutics. The improved biodistribution has the potential to safely and efficiently clear plaques from deeper brain regions via a ubiquitous vascular delivery mechanism that is scalable to larger human brains, as well as reduce the dose required to achieve the same quantity of plaque removal. Aβ is a target that uniquely benefits from the enriched capillary-and postcapillary venule-associated TfR transport process, allowing ATV formatted anti-amyloid antibodies to readily access to the brain parenchyma. The lack of arterial, leptomeningeal, and perivascular localization, and the elimination of ARIA-like events in a preclinical mouse model suggests ATV^cisLALA^ possesses unique properties for developing safer and more efficacious amyloid therapeutics.

## Acknowledgments

The authors would like to thank Butch Benitez, Kevin Rebadulla, and Dominic Sobrepena for providing animal care and support at Denali Therapeutics; Joseph McInnes and Amanda Sani for helpful discussions; and Grethe Skovbjerg, Urmas Roostalunand Jacob Hecksher-Sørensen from Gubra for technical services, support and advice.

## Conflict of Interest

MEP, NK, WK, SLD, CBD, TE, DJ, ER, DC, JCD, KG, RM, IB, RC, JC, AJC, MSD, JD, LF, JAG, MSK, DJK, AWL, HNN, ERT, PES, LS, APS, HS, RT, MEC, RJW, RGT, JL, and YZ are currently or were previously paid employees of Denali Therapeutics Inc. EDP, JA, ML, SH, JS, ACSA, PHW, DMW, and TB are currently or were previously paid employees of Biogen. Denali has filed patent application no. PCT/US2019/012990 and PCT/US2024/021179, each of which are related to the subject matter of this paper. MSD, MSK, WK, APS, and YZ are inventors of PCT/US2019/012990. KG, NK, MEP, and YZ are inventors of PCT/US2024/021179.

## Author Contributions

**Conceptualization.** MEP, EDP, NK, WK, JA, SLD, PES, MLB, APS, MEC, RJW, RGT, PHW, DMW, JL, TB, YZ

**Formal analysis:** MEP, EDP, NK, WK, JA, SLD, DJ, RM, MEC

**Investigation:** MEP, EDP, NK, WK, JA, SLD, CBD, TE, DJ, MLB, ER, DC, JCD, KG, RM, JS, ACSA, IB, RC, JC, AJC, JD, LF, JAG, DJK, AL, HNN, ERT, LS, HS, RT, APS, MEC, MSK

**Writing – original draft:** MEP, EDP, NK, WK, MEC, RGT, TB, YZ

**Writing – reviewing and editing:** MEP, EDP, NK, CBD, DJ, MEC, RJW, RGT, PHW, DMW, JL, TB, YZ

**Supervision.** EDP, KG, RM, MSD, PES, APS, HS, MEC, RJW, PHW, DMW, JL, TB, YZ

## Materials and Methods

### Expression and purification of recombinant antibodies and ATVs

ATVs were generated as previously described (Kariolis et al., 2020). Briefly, three expression plasmids consisting of the heavy chain ATV-knob, heavy chain hole, and light chain were co-transfected in Expi293 or ExpiCHO cells at a ratio of 1:1:2 according to manufacturer’s instructions. Recombinant ATV variants were subsequently purified from conditioned media by loading supernatant over a protein A column (GE Mab Select SuRe). The column was washed with 10 column volumes of PBS, pH 7.4. ATVs were eluted with 50 mM sodium citrate, pH 3.0 containing 150 mM NaCl, and immediately neutralized with 200 mM arginine, 137 mM succinic acid, pH 5.0. The ATVs were further purified by size-exclusion chromatography (GE Superdex200) using 200 mM arginine, 137 mM succinic acid, pH 5.0 as running buffer. The purified ATV molecules were confirmed by intact mass LC/MS, and purity of >95% was confirmed by SDS-PAGE and analytical HPLC-SEC.

### TfR affinity measurements by surface plasmon resonance

The affinities of TfR-binding molecules for recombinant human human TfR^apical^ were determined by SPR using a Biacore™ 8K instrument in 1X HBS-EP+ running buffer (GE Healthcare Life Sciences, BR100826). Molecules were immobilized on a Series S CM5 chip (Cytiva, 29149603) using a Human Fab Capture Kit (Cytiva, 29234601) at 10 μg/mL at a flow rate of 10 μL/min for 30 s. Serial 3-fold dilutions of recombinant human TfR^apical^ at concentrations of 24.5, 74.0, 222, 667, and 2000 nM were injected at a flow rate of 30 μL/min for 60 s followed by a 60 s dissociation using a single cycle kinetics method.

For affinity evaluation of binding to full-length human TfR ECDs, a Biacore series S sensor chip SA (Cytiva, 29104992) was immobilized with recombinant human TfR ECD at 1 μg/mL at a flow rate of 10 μL/min for 30 s. Serial 3-fold dilutions of the molecules at concentrations of 24.5, 74.0, 222, 667, and 2000 nM were injected at a flow rate of 30 μL/min for 60 s followed by a 120 s dissociation.

Data analysis was conducted using Biacore Insight Evaluation software (version 2.0.15.12933). 1:1 Languir model of simultaneous fitting of K_on_ and K_off_ was used for human TfR binding kinetics and affinity analysis.

The approximate binding affinity of TfR-binding molecules to human TfR are listed in the figure legends for each experiment.

### ADCC and CDC assays

For ADCC assays, target cells (Ramos or A20) were plated at 10,000 cells/well and opsonized with antibodies or ATVs of interest for 30 min at 37°C. Effector natural killer (NK) cells were isolated from human peripheral blood using Rosette Sep NK isolation kit (Stem Cell Technologies, Vancouver, Canada). NK cells were stimulated overnight with IL-21 (20 ng/mL, R&D Systems, Minneapolis, MN). Effector cells were added to target cells at 25:1 effector:target cells ratio (250,000 cells/well) for 4 h. Cytotoxicity was evaluated by LDH expression using CytoTox 96 Non-radioactive cytotoxicity assay kit (Promega, Madison, WI). Results were normalized to the control without any polypeptides and calculated as the percentage of maximum lysis (1% Triton X-100) in target cells.

For CDC assays, target cells (CHO-hTfR or Raji) were plated at 200,000 cells/well in serum-free media. Briefly, CHO cells were engineered to stably overexpress TfR. As above, cells were opsonized with antibodies or ATV variants for 30 min at 37°C. 50 mL of diluted baby rabbit complement proteins (Bio-rad Laboratories, Hercules, CA) was added to each well and cells were incubated for 4 h. Cytotoxicity was evaluated by LDH expression using the CytoTox-ONE homogeneous membrane integrity assay kit (Promega, Madison, WI). Results were normalized to the control without antibody or ATV and calculated as the percentage of maximum lysis in target cells.

### Mouse care and use

All procedures in animals were performed in adherence to ethical regulations and protocols approved by Denali Therapeutic or Biogen Institutional Animal Care and Use Committee. Mice were group-housed when possible, provided enrichment, and had access to water and standard rodent diet *ad libitum* on a 12-hour light/dark cycle. Age and sex of animals were evenly distributed across treatment groups and time points where possible. Before and during studies mice were monitored for health issues and excluded from study inclusion or downstream analyses if serious concerns were observed. Animals were pseudo-randomized into experimental groups with cage-mates distributed into unique experimental groups when possible. Animals and samples were assigned a unique ID to pseudo-blind and randomize samples during analysis.

### FAM (fluorescein)-labelled Aβ_1-42_ fibrils

FAM-labeled Aβ_1-42_ (0.5 mg) (Anaspec AS-23525-05) was resuspended with 100 μL DMSO followed by 1 mL PBS dilution to 100 μM. The solution is incubated at 37°C with shaking for 24 h. The FAM-amyloid beta fibrils was then transferred to a 1.5 mL ultracentrifuge tube and spun at 100,000 *g* for 30 min at 4°C in an ultracentrifuge. The supernatant was then discarded, and the pellet was resuspended in 1 mL PBS followed by extensive pipetting. The ultracentrifugation was then repeated followed by two additional rounds of washing. The pellet was resuspended in a final volume of 111 μL PBS and stored at -80°C. Dynamic light scattering (DLS) was used to characterized Aβ preparations.

### Dynamic Light Scattering (DLS)

30 μl of Aβ solution (1 mg/ml) was loaded into a 384-well plate black with clear bottom (Costar). The plate was sealed with a film and centrifuged at 1000 rpm for 5 min and loaded onto DynaPro Plate Reader III (Wyatt Technology). Readings were performed using the Dynamics v7 software at 25°C with a 5 s reading per well and an average of 10 measurements.

### *Ex vivo* phagocytosis

Naïve TfR^mu/hu^ KI (3 months old, males) were perfused with 1x PBS and the brains (including cerebellum excluding the olfactory bulb) were used for single cell dissociation. Cells were dissociated according to kit instructions using the Adult Brain Dissociation Kit (Miltenyi Biotec, 130-107-677) and gentleMACS™ Octo Dissociator. To eliminate excessive debris and myelin, the kit’s Debris Removal Solution was applied, and the final cell pellet was resuspended in 200 uL 0.5% BSA in dPBS (with calcium and magnesium). To quantify the number of live microglia per sample, a small fraction of the cells was taken and stained with Cd11b-BV421 (BioLegend 101251, 1:100), CD45-APC (BD Biosciences Cat 559864, 1:100), and Fc block (BioLegend 101320, 1:100) for 15 min at 4°C followed by washes and resuspension in FACS buffer (1% BSA + 1 mM EDTA in PBS) containing propidium iodide (PI; Miltenyi Biotec, 130-093-233). This cell fraction was then mixed with CountBright Plus Absolute Counting Beads (Invitrogen, RefC36995) and loaded on an FACSAria III Cell sorter (BD Biosciences). By using the number of live microglial cells calculated from this fraction, cells from each sample were then split into the respective treated samples such that each contained 50,000 live microglia combined with 10 nM of the respective antibody and 100 nM FAM-Aβ_1-42_ fibrils. The treated samples were then incubated at 37°C for 30 min with gentle agitation every 15 min. Control samples were treated with 10 nM of the respective antibody and 100 nM FAM-Aβ_1-42_ fibrils but incubated at 4°C instead for 30 min. Following these incubations, the treated samples were washed and stained with Cd11b-BV421 (BioLegend 101251, 1:100), CD45-APC (BD Pharmingen, 1:100), and Fc block (BioLegend 101320, 1:100) for 15 minutes at 4°C followed by washes and resuspension in FACS buffer containing PI. On the sorter, 10,000 live microglia were recorded per treated sample along with the intensity of the FAM signal. To quantify the intensity of FAM signal per live microglial cells analysis was performed on FlowJo 10.8.1.

### PKPD and tissue harvest studies in TfR^mu/hu^ KI mice

TfR^mu/hu^ KI mice (Kariolis et al., 2020) or 5xFAD; TfR^mu/hu^ KI mice were restrained and received a single intravenous tail vein injection of anti-Aβ or ATV:Aβ molecules. In-life blood was collected by submandibular or submental puncture with a lancet into EDTA coated tubes (Sarstedt Microvette 500 K3E, 201341102). At terminal time points, mice were deeply anesthetized with 2.5% avertin (intraperitoneal, IP); terminal blood was collected by intracardiac puncture and collection into EDTA coated tubes. If collected, CSF was then drawn by making a skin incision to expose the neck muscles, making a blind puncture into the cistern magna with the tip of a needle, and collecting CSF into a pulled glass capillary tube before ejecting CSF into a low-protein-binding tube. Plasma and final CSF samples were generated following centrifugation of drawn blood or CSF at 18,213 *g* for 7 minutes at 4°C. Mice were then perfused transcardially with ice cold PBS for approximately 4-6 min at 5 mL/min using a peristaltic pump (Gilson, Minipuls 3). Freshly perfused and extracted brains were dissected on a cold block on wet ice and divided for various endpoints, either immediately snap-freezing in tubes on dry ice or being immediately immersed in 4% paraformaldehyde (Electron Microscopy Sciences, 15714-S) diluted in PBS for fixation overnight at 4°C. If desired, whole blood was collected from animals that did not receive in-life bleeds into EDTA tubes and kept at 4°C for quantification of reticulocytes by a veterinary lab (Quality Vet Lab, Davis, CA, USA).

### Bone marrow flow cytometry

Following perfusion of the mice, femurs and/or tibias were collected and all muscle removed, the epiphysis cut from the shaft of the bones, and placed into a 0.5 mL tube with a small hole at the bottom inside a 1.5 mL tube, and the marrow centrifuged out fresh bone marrow at 18,213 *g* speed for 3 min at 4°C.

Bone marrow samples from each animal were dissociated into single cells in 200 μL of FACS buffer (2% FBS and 1mM EDTA in PBS). After spun down at 300 g for 5 min in FACS tubes, the supernatant was discarded, and the cells were resuspended in 200uL of FACS buffer. 40 μL of sample and 10 μL of control were added to a 96-well, round-bottom plate. The plate was washed with 100 μL of FACS buffer per well and spun down at 300 *g* for 5 min. After emptying the plate, the cells were resuspended and blocked with 100 μL per well of 1:50 Mouse-Fc block (BioLegend #101320) in FACS buffer for 5 min. The plate was spun down at 300 *g* for 5 min before staining with 100 μL per well of Fixable Viability Dye (BD Biosciences #564406), mouse anti-CD71/APC, rat anti-CD44/APC, rat anti-TER119/PE, and isotype control antibodies for 30 min. The plate was washed, spun down, and emptied two more times before the cells were resuspended in 75 μL FACS buffer per well.

Samples were analyzed on a FACS Canto II (BD Biosciences). To isolate the population of CD44+ reticulocytes, gates were applied in the order of cells (SSC-A/FSC-A), single cells (FSC-H/FSC-A), live cells (SSC-A/Am-Cyan-A), TER119+ (SSC-A/PE-A), and erythroids (APC-A/FSC-A). The populations of reticulocytes lie in APC-A+ and 0∼50K of FSC-A. CD44+ reticulocytes were analyzed and quantified in FlowJo. Percentage of CD44+ in TER119+ were exported and plotted in GraphPad Prism.

### Acute multidose phagocytosis study in 5xFAD; TfR^mu/hu^ KI mice

5xFAD; TfR^mu/hu^ KI mice (mixed sex, *n*=11-12 per group) received IP injections of anti-Aβ, ATV^LALA^:Aβ, or ATV^cisLALA^:Aβ (approximate affinity for huTfR apical domain of 500 nM) on days 0, 3, 6, and 9 and were perfused as described above and brains collected for immersion-fixation (24 in 4% PFA) or flash-freezing on day 12. Hemibrains were stored at 4°C in 15% sucrose then transferred to 30% sucrose in PBS until the brains sunk fully. Cryoprotected brains were sectioned on a freezing microtome (40 μm, sagittal) and sections stored in PBS with 0.01% sodium azide prior to immunohistochemistry.

Four brain sections per animal were selected for microglia and plaque immunohistochemistry; two sections were approximately 3 mm lateral to the midline and two sections were approximately at the midline (0 mm lateral). Free-floating sections were incubated for 2 h in blocking solution (5% donkey serum in PBS with 0.3% triton X-100) at room temperature (RT), incubated overnight in blocking solution containing primary antibodies at 4°C, washed 3x for 15 min each in PBS with 0.3% triton X-100, incubated 2 h at RT in blocking solution containing secondary antibodies, incubated 20 min in DAPI (Thermo, D1306) solution in PBS with 0.3% triton X-100, and washed 3x for 15 min each in PBS with 0.3% triton X-100. Sections were mounted onto slides and coverslipped using Prolong Glass Antifade mounting medium. Primary antibodies included: rabbit-anti-human amyloid β (IBL America, #18584; 1:500), rat-anti-CD68 (BioRad, MCA1957; 1:500 and secondary antibodies included: donkey-anti-rabbit IgG AF488 (Life, A21206; 1:500), cross-adsorbed donkey-anti-rat IgG Dy555 (Invitrogen, SA5-10027; 1:500).

Three brain sections per animal were selected for plaque and human IgG immunohistochemistry; sections were approximately 3 mm lateral to the midline. Free-floating sections were incubated for 1 h in blocking solution (5% donkey serum in PBS with 0.3% triton X-100), incubated 2 h in blocking solution containing rabbit-anti-human amyloid beta (IBL America, #18584; 1:500) and donkey-anti-human AF647 (Jackson ImmunoResearch, 709-606-149; 1:250), washed 3x for 15 min each in PBS with 0.3% triton X-100, incubated 1 h in blocking solution containing donkey-anti-rabbit AF488 (Life, A21206; 1:500) and incubated 20 minutes in DAPI solution in PBS with 0.3% triton X-100, and washed 3x for 15 min each in PBS with 0.3% triton X-100. All steps were performed at room temperature. Sections were mounted onto slides and coverslipped using Prolong Glass Antifade mounting medium.

Slides were imaged using a Zeiss Axioscan.Z1 slide scanner at 20X magnification and images were processed using custom macros in Zeiss ZEN software. Image processing macros automated the following steps for all images: median smoothing, extracting objects and channels of interest, channel processing (e.g., rolling-ball background subtraction, threshold equally across all animals to identify signal eliminate background, set minimum and maximum size cutoff to eliminate noise/debris). This created additional binary image channels for each processing step. The ZEN software was then used to analyze the processed images by first auto-segmenting tissue from background using DAPI channel image (smoothing and filling holes) to determine tissue section area. Binary image channels were then analyzed to measure the plaque area (by size), area of CD68+ microglia and overlap with plaques and their immediate surroundings (‘dilated’ plaques). Plaque size bins (30-125 μm^2^, 125-250 μm^2^, 250-500 μm^2^, 500-3,300 μm^2^) were selected based on a previously published study (Sevigny et al., 2016).

### Biodistribution in normal TfR^mu/hu^ KI mice

Brains were transferred to PBS + 0.01% sodium azide following immersion fixation and shipped to Neuroscience Associates for sectioning using MultiBrain Technology; up to 20 hemibrains were embedded into a single gelatin block and sectioned in the sagittal plane (35 μm). Sections were collected in a series of 12 and stored in antigen preservation solution (50% PBS:50% ethlyene glycol + 1% PVP) at -20C until staining. Three to four sections per animal were allowed to come to room temperature and then washed in PBS before incubation in blocking buffer (PBS/TBS + 1% BSA + 1x fish gelatin (BioWorld 21761058+ 0.5% triton X-100 + 0.1% sodium azide) for 3 hours at room temperature on a rocker. Sections were then incubated overnight in antibodies (NeuN, Abcam, ab1777487, 1:1000; CD31, R&D Systems, Af3628, 1:500; and donkey F(ab’)2-anti-huIgG, Jackson, 709-606-149, 1:250) at 4°C, followed by 3x 15 min washes and incubation with secondary antibodies (1:500; donkey-anti-rabbit, Invitrogen, A11042; donkey-anti-goat, Invitrogen, A11055) and DAPI (5 μg/mL,Thermo, D1306), before 3x 15 min washes, mounting, and coverslipping with Prolong Glass (Invitrogen, P36984).

### Target engagement in 5xFAD; TfR^mu/hu^ KI mice

Brains were transferred to PBS + 0.01% sodium azide following immersion fixation and shipped to Neuroscience Associates for sectioning using MultiBrain Technology; up to 20 hemibrains were embedded into a single gelatin block and sectioned in the sagittal plane (40 μm). Every tenth section (at an interval of 400 μm) was stained free-floating at Neuroscience Associates. All incubation solutions in following steps used Tris buffered saline (TBS) with Triton X-100 as the vehicle; all rinses were with TBS.

After rinses, the sections were treated in formic acid rinsed and stained with rabbit-anti-human amyloid β (IBL America, #18584; 1:500) and donkey F(ab’)_2_-anti-huIgG (Jackson, 709-606-149, 1:500), overnight at room temperature. Following rinses, secondary antibody (donkey-anti-rabbit AF647, Jackson ImmunoResearch 711-605-152, 1:500) was applied. The sections were again rinsed, then treated with a Hoechst counterstain. Following further rinses, the section was mounted on gelatin coated glass slide, then air dried. The stained slide was dehydrated in alcohols, cleared in xylene and coverslipped. Slides were imaged and analyzed as described above using a Zeiss Axioscan.Z1 slide scanner at 20X magnification and custom macros to identify plaques and huIgG overlap with plaques.

### iDISCO brain clearing

5xFAD; TfR^mu/hu^ KI mice (female, 3-3.5 months, *n*=7 per group) received a single 10 mg/kg IV injection of AF750-conjugated-anti-Aβ or AF750-conjugated-ATV^cisLALA^:Aβ (approximate affinity for huTfR apical domain of 500 nM). 24 h after dosing mice were deeply anesthetized and transcardially perfused with ice cold heparinized PBS (15 U/mL) for approximately 5-6 min followed by approximately 5-6 min of RT 10% neutral buffered formalin (NBF), both at 5 mL/min using a peristaltic pump (Gilson, Minipuls 3). Brains were post-fixed in NBF at RT for 24 h, rinsed in PBS, then transferred to PBS + 0.02% sodium azide and stored at 4°C until staining.

For one hemisphere, *n*=2 animals per group were stained with a secondary for the huIgG backbone of the dosed molecules (**Fig4l-m)** and *n*=5 animals per group were stained with Congo red for detection of plaques and direct detection of the AF750 conjugate. For the other hemisphere, *n*=1 animal per group was used for pilot staining and the remaining *n=6* animals per group were stained with αSMA, Aβ, and huIgG (**Fig 4o-p**, **Supplementary Fig3**).

For staining, hemi-brains were dehydrated in a methanol/water gradient on a gentle rocker, during which they were incubated for an hour each in 20%, 40%, 60% 80%, and 100% methanol (1-2x), followed by an overnight incubation in 66%dichloromethane (DCM)/33% methanol at RT. Samples were then washed twice in 100% methanol, chilled in fresh 100% methanol and bleached in fresh-chilled 5% H_2_O_2_ in methanol overnight at 4°C. After bleaching, samples were rehydrated in a methanol/PBS + 0.2%Triton-X100 series of 80%, 60%, 40%, and 20% methanol at RT, after which they were washed in PBS+ 0.2%Triton-X100 and permeabilized for 3 days at 37°C in a PBS+ 0.2%Triton-X100 buffer made with 20% DMSO, 0.02% glycine and 0.02% sodium azide. Blocking and staining steps were then carried out according to the details below for either the Congo red or huIgG, αSMA, and Aβ staining sets, or for the parallel experiment elucidating vascular anatomy, SM22. Samples were then cleared in a methanol/water gradient at RT on a gentle rocker (20%, 40%, 60% 80%, and 100% methanol), incubated in 100% methanol overnight at RT, washed in 100% methanol once more and then incubated in 66% Dichloromethane (DCM)/33% methanol for 3 hours at RT. Samples were then washed in 100% DCM twice for 15 min before index-matching in Dibenzyl ether (DBE). Congo Red stained samples were imaged in DBE, while huIgG, aSMA, Aβ stained samples were switched to ethyl cinnamate (ECi) at least 24 hours before imaging in ECi.

### Congo Red (*n*=5/group) or huIgG staining (*n*=2/group), or SM22 (*n*=6/group)

After permeabilization, samples were blocked for two days (37°C) in PBS with 0.2%Triton-X100, 10% DMSO, 6% normal donkey serum, and 0.02% sodium azide. Samples were incubated with Congo Red (15 μM, Millipore Sigma, BioXtra, C6277) or donkey-anti-huIgG-Cy5 (1:250, Jackson ImmunoResearch, 709-175-149) in PTwH (PBS with 0.2% Tween-20 and 5mg/L heparin, H3149-50KU, EMD Millipore))/3% donkey serum at 37°C for 7 days and then washed in PTwH. For the parallel experiment elucidating arterial anatomy, blocked brains were incubated in anti-SM22 primary antibody (1:1000, Abcam, ab14106) diluted in PBSG-T and 0.1% saponin (10 μg/mL) for 14 days with agitation at room temperature. Following primary antibody incubation, the samples are washed 3 x 2 h in PBST, left over two days in PBST. The secondary antibody incubation was carried out for 7 days in donkey anti-rabbit AlexaFluor488 (1:500, Jackson ImmunoResearch, 711-545-152) diluted in PBSG-T and 0.1% saponin (10 μg/mL), with agitation at room temperature. Following secondary antibody incubation, the samples are washed 3 x 2 h in PBST and incubated in PBST over 3 days.

### huIgG, αSMA, Aβ staining (*n*=6/group)

After permeabilization, samples were blocked for three days (37°C) in PBS with 0.2%Triton-X100, 10% DMSO, 6% normal donkey serum, and 0.02% sodium azide. Blocked brains were incubated in anti-huIgG-DL800 (1:500, Invitrogen, SA510132), anti-Aβ (1:400, IBL America, #18584), and anti-αSMA-DL550 (1:500, Novus, NBP2-34522R) at RT for 2 weeks on gentle rocking speed. All antibodies were diluted in a PBS-based antibody incubation buffer containing 0.2% gelatin, 0.5% tritonX100, and 0.1% saponin. Following incubation, tissue was washed with PTwH for 1×10 min, 1×20 min, 1×30 min, 1x 1 hr, 1x 2 hr at RT and 1×3 days at 4°C before incubation in secondary antibody. The secondary antibody incubation was carried out in antibody incubation buffer (with an additional 5 mg/L heparin) for 7 days at RT, with donkey anti-rabbit Cy5 (1:500, Jackson ImmunoResearch, 711-175-152), and then washed in PTwH.

### Whole hemibrain imaging and quantification

iDISCO cleared brains were imaged on a Lavision UltraMicroscope (Lavision Biotec GmbH) with an MVPLAPO 2X objective/0.5 NA with a 0.63x zoom. Hemibrains were imaged to collect autofluorescence and relevant channels. Additional imaging was performed using a Miltenyi Blaze (Miltenyi Biotec) light sheet microscope using a 1.1x/0.1 NA MI PLAN objective with a 1.0x zoom (**Supplementary Fig 3**). Fully cleared samples were imaged in the Miltenyi Blaze with both the 1x objective (1.1x zoom, 0.09 numerical aperture, fixed focus and blending) and the 12x objective (1x zoom, 0.19 numerical aperture, adaptive focus and adaptive blending). All images per magnification were taken with the same optimized settings with bidirectional illumination. 1x images were taken at 60% sheetwidth, at 2.4 μM steps. Excitation (Emission) as follows: 785 (805): 15%, 640 (680) 10%, and 561 (620) 10%. 12x images were taken at 10% sheetwidth, at 0.54 μM stepsize. Excitation (Emission) as follows: 785 (805): 25%, 640 (680) 9%, and 561 (620) 8%. Brain volumes were visualized in Imaris v9.9. Plaques and vasculature were segmented using custom code developed in python v3.9 using the following pipeline. First, a tissue mask was extracted from the volume by thresholding the αSMA image at a level that also segmented the autofluorescence followed by Boolean operations to fill holes and remove small objects using the morphology module of scikit-image v0.18 (van der Walt et al., 2014). Inhomogeneity in the Aβ and huIgG illumination fields was estimated by smoothing each channel with a gaussian filter, subtracting the smoothed image from the original image, and then clamping all negative intensity values to zero. Next, plaques were segmented by thresholding the illumination corrected Aβ image followed by separation of individual plaques using the label function in scipy v1.7 (Virtanen et al., 2020). Major arteries and arterioles were segmented by thresholding the αSMA at a higher level that excluded the autofluorescence followed by Boolean operations to remove small segments and holes. Imaris was used to apply the plaque and artery masks to the huIgG channel to visualize the co-localization. Intensity of huIgG and Aβ staining were found to vary with depth from tissue surface, likely a technical artifact of the clearing and staining process in this experiment, but co-localization of huIgG signal with both Aβ and αSMA via a Pearson’s correlation coefficient (PCC) was less impacted by this gradient effect. Mean intensity of huIgG, Aβ, and αSMA as well as the Pearson’s correlation coefficient between Aβ and huIgG and between αSMA and huIgG were calculated per plaque and then a volume-weighted average was calculated per animal. Two tailed t-tests were performed using the ttest_ind function from scipy’s stats module. To preserve normality assumptions, correlation coefficients were first transformed to z-scores using Fisher’s z-transformation before averaging or calculating t-tests.

### Brain homogenization and huIgG ELISA

Fresh-frozen brain samples were homogenized in 10x volume/weight of lysis buffer (1% NP40 (Thermo 85124) in ice cold PBS with protease and phosphatase inhibitors (Roche 04693132001 and 04906837001)) using a 3 mm bead (Qiagen 69997) and agitation with the Tissue Lyser II (Qiagen, 85300). Samples were homogenized by two 3 min sessions at 27 Hz at 4°C. Following incubation on ice for 20 minutes samples were centrifuged for 20 minutes at 14,000 *g* at 4°C. The supernatant was removed and stored at -80°C until analysis. As this method does not completely solubilize Aβ, it is expected that much of the huIgG is not recovered in the supernatant and represents a free or soluble fraction of the total huIgG.

Concentration of huIgG in plasma and brain lysates were quantified using a generic anti-human IgG sandwich-format ELISA as previously described (Chew et al., 2023). Plates were coated overnight with donkey anti-human IgG (JIR #709-006-098) at 1 μg/mL in sodium bicarbonate solution (Sigma #C3041-50CAP) with gentle agitation at 4 °C. Plates were then washed with wash buffer (PBS + 0.05% Tween 20) three times. Assay standards and samples were diluted in PBS + 0.05% Tween 20 and 1% BSA. Standard curve was prepared at a range from 0.41 to 1500 ng/mL or 0.003 to 10 nM (BLQ< 0.03 nM). Standards and diluted samples were incubated for 2 h with agitation at RT. After incubation, plates were washed with wash buffer three times. The detection antibody, Goat anti-human IgG (Jackson ImmunoResearch, 109-036-098) was used as the detection antibody and was diluted in blocking buffer (PBS + 0.05% Tween-20 + 5% BSA) to a final concentration of 0.02 μg/mL. Plates were incubated for 1 h with agitation at RT, washed 3x, and then developed by adding TMB substrate and incubated for 5–10 min followed by quenching with 4N H2SO4; plates were read using 450 nm absorbance.

### ARIA animal model

5xFAD; TfR^mu/hu^ KI mice were 11-13 months of age at the start of the studies. All experimental procedures were performed in accordance with the standards of Biogen IACUC and the Cambridge Laboratory Animals Ordinance, Cambridge MA (Ordinance 1086).

### ARIA experimental study design

The objective for the first ARIA study was to assess the potential for reduced ARIA incidence upon treatment with ATV:Aβ and for the second ARIA study was to assess the role of effector function and ATV-mediated brain delivery in the potential reduction of ARIA incidence. Following the collection of a baseline MRI scan, 5xFAD;TfR^mu/hu^KI mice were administered weekly IP injections of the treatments for 10-12 weeks, and MRI imaging was performed within 1-3 days after weekly dosing. Untreated animals were used as control and were scanned at baseline and at end of study. Males were used for the first study and mixed sex (equally distributed between groups) were used for the second study. Anti-CD4 antibody (Ref BP0003-R050, BioXCell) was administered IP every 2 weeks, starting 1 day before the first dosing of compounds (0.5 mg/animal). In the second study, blood was collected from all animals prior to the first dose and prior to the last dose, and in both studies blood was collected at termination. Blood was processed to plasma and stored at -80C for analysis.

### Preparation of brain tissue for ARIA studies

Animals were euthanized with CO_2_ asphyxiation 24 h after the last dose, and blood was collected by cardiac puncture. Cardiac perfusion was initiated with cold PBS pH 7.2 (Gibco, Ref 2645342), followed by 20 ml of cold 10% NBF for fixation of the meninges. The whole brain with the meninges attached was dissected out of the skull and drop fixed in 10% NBF STS containers for 1-2 days (Epredia, Ref 53601).

### ARIA MRI protocol

Mice were anesthetized with isoflurane in medical oxygen (2% induction, 1-2% maintenance). Images were acquired on a 9.4T preclinical MRI (Bruker BioSpec 94/20, Billerica, MA, USA) using Paravision (6.0.1) with a mouse head RF coil. Mice were secured into the head coil using a bite bar, and respiration was kept between 70-160 beats per minute. Body temperature was maintained at 37 ± 0.3 °C using an MR compatible warm air heater system (Small Animal Instruments, Inc. Stony Brook, NY, USA), and temperature recorded using a rectal probe. Baseline scans were first acquired to document the absence of pre-treatment abnormalities. Weekly scans were acquired as follows: we first acquired T2 scan (slice thickness= 0.5mm, repetition time [TR]= 4100ms, echo time [TE]=60 ms, averages=10, repetitions=1, echo spacing=7.5ms, rare factor=16, flip angle=90° FOV=16×16mm, matrix= 128×128, slices=30), followed by T2* MGE (slice thickness=0.5mm, TR=1800ms, TE=3ms, averages=2, repetitions=1, echo images=12, echo spacing=4.5ms, minimum echo spacing=1.58ms, flip angle=30°, FOV=16×16mm, matrix= 160×160, slices=30), and DWI (slice thickness=0.5mm, TR=3200m, TE=21.5ms, averages=1, repetitions=1, segments=2, flip angle=90°, FOV= 16×16mm, matrix=128×60, slices=30).

### ARIA-related pathology assays and analysis

Whole brains were processed following post-fixation in 10% neutral-buffered formalin for 48-72 hours. The right hemisphere was gently marked with red ink. Brains were then sectioned in the coronal plane using a mouse brain block to produce a 6-piece coronal trim. The first cut was localized at the caudal-most visible plane of the optic tracts. One cut was placed rostral to the optic tract section and 3 cuts were placed caudal to the optic tract section, all at 2-mm intervals. Six coronal slabs were processed and embedded in paraffin blocks for sectioning in the rostral-to-caudal direction. Blocks were gently faced and sectioned at 5-micron thickness to produce 10 consecutive unstained slides.

One slide was stained with hematoxylin and eosin (H&E), coverslipped on a Tissue-Tak Glas automated glass coverslipper (Sakura Finetek) and imaged on a Pannoramic P250 slide scanner with a 20x objective. H&E slides were examined by EDP who was blinded to the treatment group assignments and to the MRI image data of all animals. The following grading system was applied to leptomeningeal arterioles in the 6 brain sections: Grade 0 (none) – no obvious meningovascular inflammation; Grade 1 (minimal) - perivascular mononuclear cells without obvious phagocytic morphology – when present, a focal finding; Grade 2 (mild) – subtle vascular macrophage infiltrates requiring high power examination to visualize; Grade 3 (moderate) - multiple layers of vascular macrophages that were evident at low-to-medium power; Grade 4 (severe) – any vascular inflammatory infiltrate with clear fibrinoid change. Meningovascular inflammation grade scores were 1) summarized on a per animal basis as the predominant meningovascular inflammation grade (both studies) and 2) summated for all foci of meningovascular inflammation throughout 6 brain sections for a summated meningovascular inflammation score (second study only). H&E slides were also examined for microinfarcts and hemorrhages.

One slide was stained with the Perls iron method (Prussian Blue Iron Kit; Polysciences Inc.; Cat # 24199), coverslipped and imaged on a Pannoramic P250 slide scanner with a 20x objective. Perls iron slides were examined for microhemorrhages. Leptomeningeal vascular profiles demonstrating 3 or more Perls-reactive hemosiderin foci and that were confirmed on H&E to contain hemosiderin pigment were counted (leptomeningeal hemosiderin foci). Perivascular hemosiderin foci around penetrating cortical arterioles (penetrating perivascular hemosiderin) were added to the total hemosiderin foci. Finally, microinfarcts that showed any hemosiderin staining were counted (hemosiderin-positive microinfarcts).

An immunohistochemistry assay for albumin was performed on a Ventana Discovery Ultra autostainer. Heat-induced antigen retrieval was first performed for 64 minutes in CC1 buffer. Sections were then incubated with rabbit anti-albumin antibody (Abcam Cat # ab192603) at a concentration of 0.5 Dg/mL. HQ-HRP amplification secondary antibody to detect and DAB to visualize. All assays were batched to prevent effects of run-to-run variability. Slides were imaged on a Panoramic P250 slide scanner with a 20x objective. Tissue section immunoreactivity was analyzed with custom-designed, threshold-based segmentation protocols using median filters and respective color deconvolution filters in Visiopharm Image Analysis Software (version 2019.12.0.6842 and subsequent updates). Cortical albumin immunoreactivity was calculated as the summed areas of cortex immunoreactive for albumin divided by the composite cortical area.

## Notes

### Summary of Updates

Minor revisions to title and abstract

