## Supplemental Figures & Table for "Engineering anti-amyloid antibodies with transferrin receptor targeting improves brain biodistribution and mitigates ARIA"

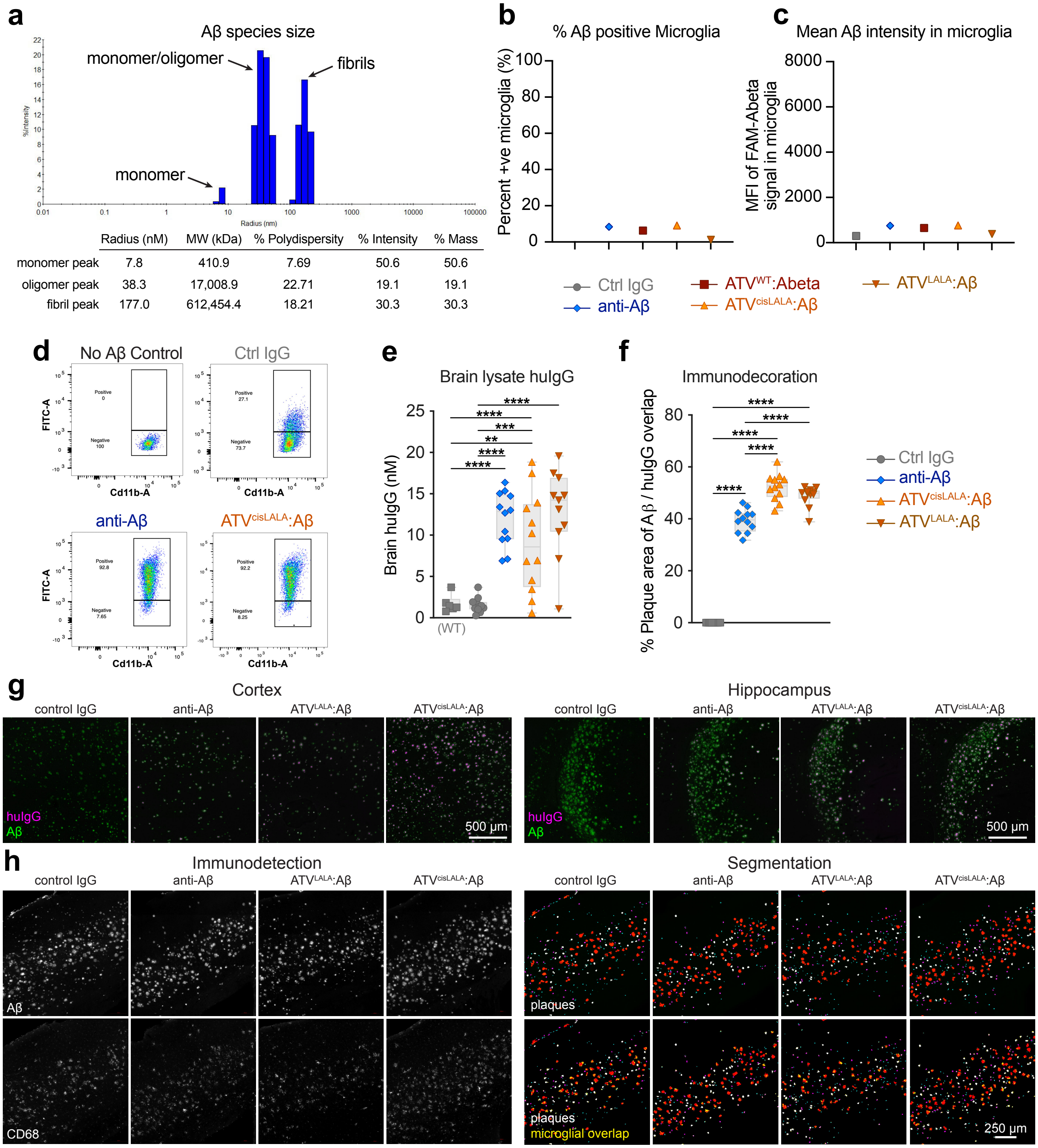


| | Radius (nM) | MW (kDa) | % Polydispersity | % Intensity | % Mass |
|---|---|---|---|---|---|
| monomer peak | 7.8 | 410.9 | 7.69 | 50.6 | 50.6 |
| oligomer peak | 38.3 | 17,008.9 | 22.71 | 19.1 | 19.1 |
| fibril peak | 177.0 | 612,454.4 | 18.21 | 30.3 | 30.3 |

**Supplementary Figure 1. Experimental controls and details for *ex vivo* and *in vivo* phagocytosis experiments. a**, FAM-labeled Aβ aggregates were characterized using dynamic light scattering (DLS); peaks for monomers, oligomers, and fibrils indicated. **b-c**, FAM-Aβ-positive live microglia (**b**) and mean fluorescence intensity (MFI) of FAM-Aβ signal in live microglia (**c**) following incubation of dissociated brain cells with FAM-labeled Aβ fibrils at 4°C. **d**, Flow cytometry diagrams and gating strategy for live microglia. **e**, Brain lysate concentrations in the cerebellum 3 days after the last dose in wild type (WT) $TfR^{mu/hu}$ KI mice, or 5xFAD; $TfR^{mu/hu}$ KI mice. **f**, Colocalized area of immunodetected Aβ and huIgG quantified from widefield whole sagittal brain sections, normalized to area of plaques. **g**, Representative images of huIgG and Aβ immunodetection from the cortex and hippocampus. **h**, Representative images of Aβ plaque signal and segmentation by size (first row; 30-125 $\mu m^2$ cyan, 125-250 $\mu m^2$ purple, 250-500 $\mu m^2$ white, 500-3300 $\mu m^2$ red) or CD68 microglial staining and microglial overlap (yellow) within and immediately surrounding plaques (second row). Only significant differences indicated on graphs; (**e**) one-way ANOVA or (**f**) two-way ANOVA with multiple comparisons; (**e**) Tukey's post hoc comparing all groups, $^{**}p < 0.01$, $^{***}p < 0.001$, $^{****}p < 0.0001$ (**f**) Tukey's post hoc comparing all groups, $^{****}p < 0.0001$. The TfR affinity for huTfR apical domain in these experiments is ~1200 nM (**b-d**) or ~500 nM (**e-h**).

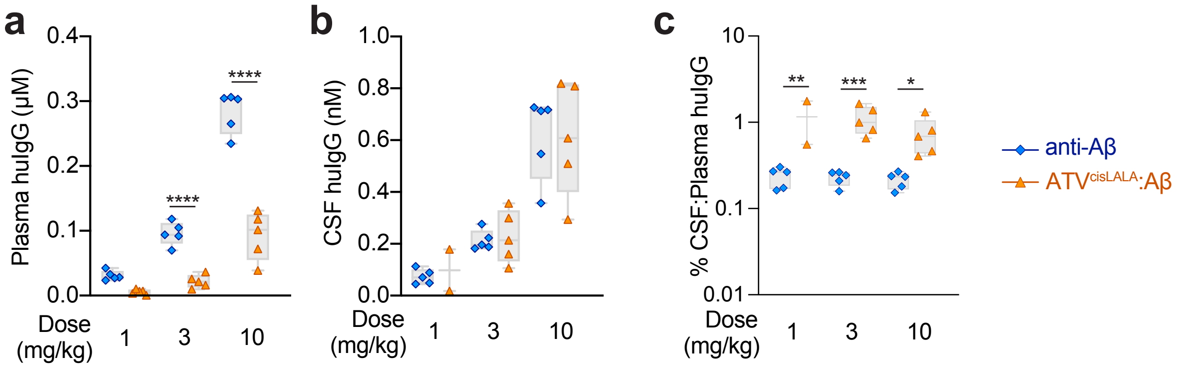


**Supplementary Figure 2. Plasma and CSF huIgG exposures. a-c,** 5xFAD; TfR$^{mu/hu}$ KI mice received a single IV dose of 1, 3, or 10 mg/kg anti-Aβ or ATV$^{cisLALA}$:Aβ. huIgG concentrations were measured by ELISA in plasma (**a**) or cisternal CSF (**b**) and the percent CSF:plasma calculated (**c**) 24 h post-dose. Only significant differences indicated on graphs; (**a-c**) one-way ANOVA; Sidak's multiple comparisons test paired by dose level, *$p$ < 0.05, **$p$ < 0.01, ***$p$ < 0.001, ****$p$ < 0.0001. The ATV affinity for huTfR apical domain in these experiments is ~500 nM.

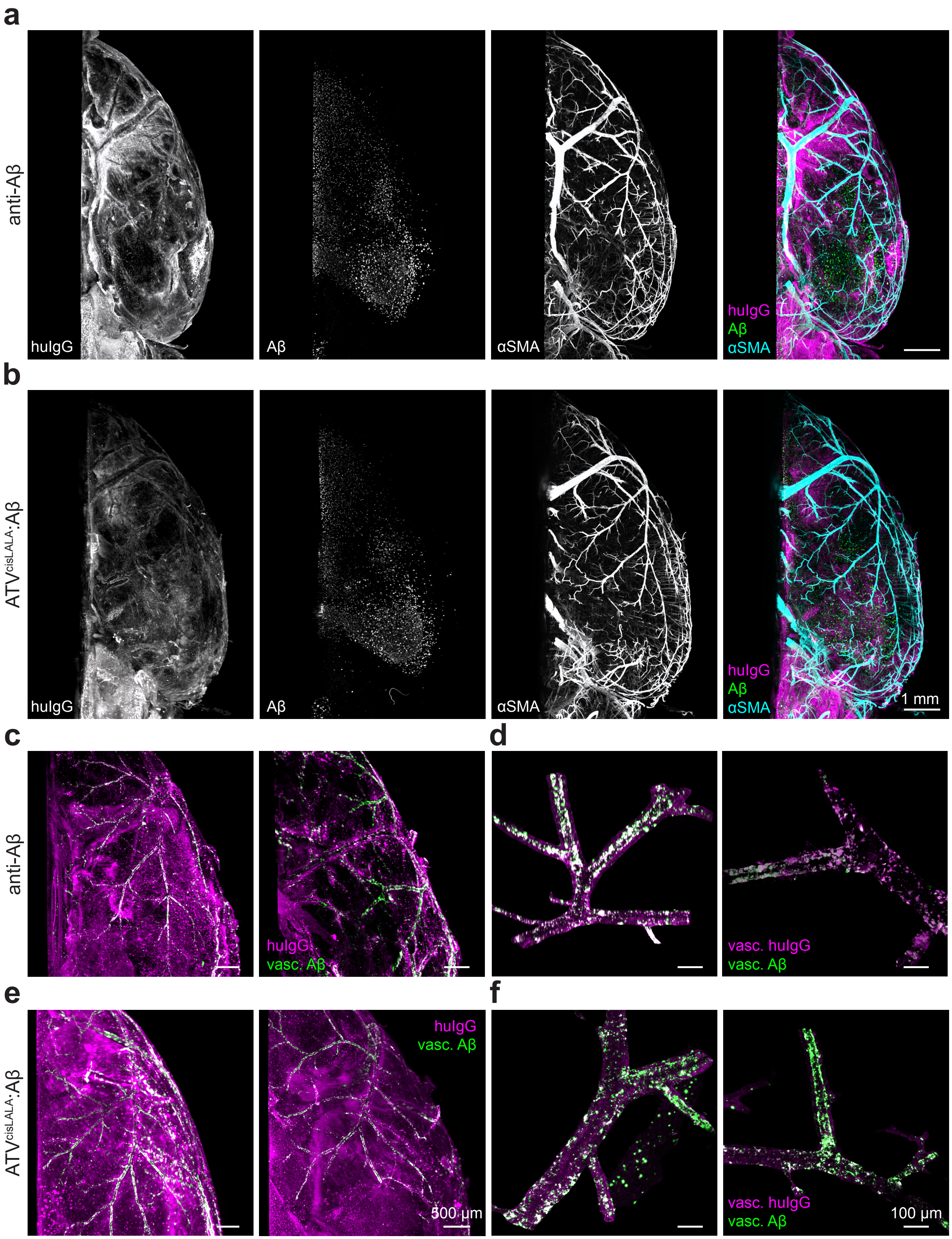


**Supplementary Figure 3. 3D hemibrain biodistribution and association with vascular amyloid. a-b**, Representative maximum intensity projection light sheet images of staining for huIgG, Aβ, and α-smooth muscle actin (αSMA) in 3-3.5 month 5xFAD; $TfR^{mu/hu}$ KI mice (*n*=6/group) after a single 10 mg/kg IV dose of AF750 conjugated anti-Aβ (**a**) or $ATV^{cisLALA}$:Aβ (**b**). **c,e,** Representative images of immunodetected huIgG and vascular amyloid (masked based on close proximity to smooth muscle cells) from two animals per group with apparent CAA. **d, f**, Higher magnification imaging of the masked vascular amyloid signal and masked vascular huIgG signal showing colocalized signal (white), if present.

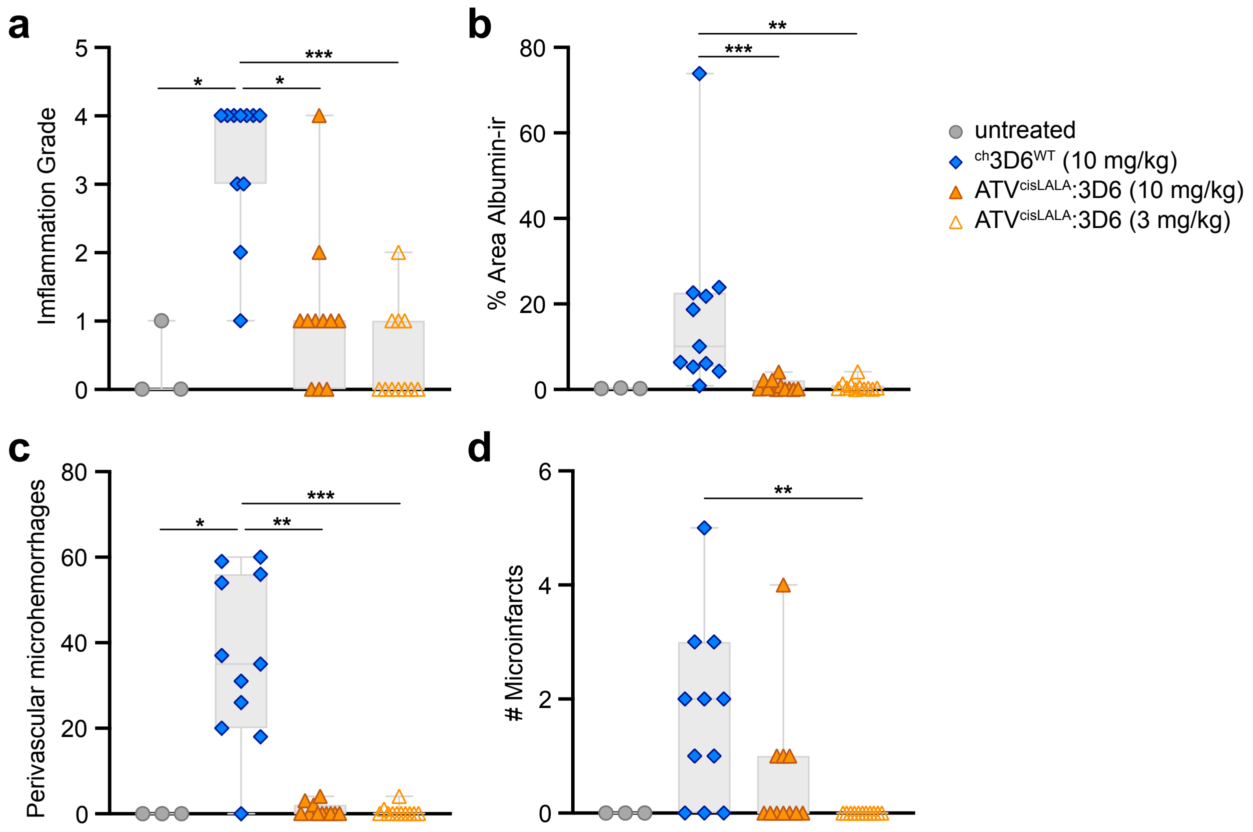


**Supplementary Figure 4. $^{ch}$3D6-induced meningovascular pathology is mitigated by both TfR-mediated brain uptake at equal or exposure-matched doses. a-d,** Blinded histopathological assessment of brains from male 5xFAD; TfR$^{mu/hu}$ KI mice treated with $^{ch}$3D6$^{WT}$ and dose-matched (closed triangle) or exposure-matched (open triangle) ATV$^{cisLALA}$:3D6 starting at 3 or 1.75 mg/kg and increasing to 5 or 10 mg/kg at the 5$^{th}$ dose. Plots of the predominant meningovascular inflammation grade observed in each animal (**a**), cortical albumin immunoreactivity (**b**), Perls-reactive perivascular hemosiderin/microhemorrhage foci (**c**) and microinfarcts (**d**). Only significant differences indicated on graphs; (**a-d**) Kruskal-Wallis tests used as D'Agostino & Pearson normality test failed for at least one group; (**c**) Dunn's post hoc comparing all groups, *$p$ < 0.05, **$p$ < 0.01, ***$p$ < 0.001, ****$p$ < 0.0001. The ATV affinity for huTfR apical domain in these experiments is ~1200 nM.

### Supplemental Table 1. Description of MRI lesions

| Treatment | Animal ID | MRI Lesions: Description | # | Size | MRI Score* |
|---|---|---|---|---|---|
| $^{ch}3D6^{WT}$ | 1 | No lesion | | | 0 |
| | 9 | Convexity hyperintense unilateral lesion *scan 6 - week 10* + Convexity hyperintense unilateral lesion *scans 13-15 - week 7* | 2 | 2 | 4 |
| | 15 | Multiple lesions, hyperintense and hypointense - diffuse and focal *all levels - weeks 3-10* | >5 | >5 | 10 |
| | 17 | Convexity hyperintense bilateral lesions *scans 4-6 - weeks 6-9* + Focal hyperintense lesion *scans 9-11 - week 9* + Convexity hyperintense bilateral lesions *scans 9-14 - week 7* | 5 | 3 | 8 |
| | 20 | Convexity hyperintense bilateral lesions *scans 3-6 and 9-12 - week 7* + Focal hyperintense lesion *scans 9-12 - week 10* | 5 | 4 | 9 |
| | 27 | Convexity hyperintense unilateral lesion *scans 9-20 - week 7-8* + 3 Focal hyperintense lesions *scans 10-11 - week 10* + Diffuse hyperintense cortical lesions *scans 16-21 - weeks 9-10* | >5 | 4 | 9 |
| | 31 | Convexity hyperintense unilateral lesion *scans 13-20 - week 10* | 1 | 3 | 4 |
| | 33 | Convexity hyperintense bilateral lesions *scans 3-8 - weeks 5-10* + Convexity unilateral hyperintense lesion *scans 10-12 - week 10* + Focal hypodense meningeal lesion *scan 9 - weeks 9-10* | 4 | 3 | 7 |
| | 39 | Focal hyperintense hippocampal lesion *scans* 14-15 - week 10 | 1 | 3 | 4 |
| | 45 | No lesion | | | 0 |
| $^{ch}3D6^{kLALA}$ | 3 | No lesion | | | 0 |
| | 4 | No lesion | | | 0 |
| | 8 | Diffuse hyperintense unilateral lesion (CA1) *scans 10-14 - week 10* | 1 | 3 | 4 |
| | 16 | Convexity hyperintense bilateral lesions *scans 4-6 - week 10* + Focal hyperintense lesion *scan 5 (posterior) - week 10* + Convexity hyperintense bilateral lesions *scans 10-20 - week 10* + Medial hyperintense diffuse lesion *scans 19-21 - week 10* | 6 | 3 | 9 |
| | 21 | No lesion | | | 0 |
| | 23 | No lesion | | | 0 |
| | 34 | No lesion | | | 0 |

| | | | |
|---|---|---|---|
| | 38 | No lesion | 0 |
| | 42 | No lesion | 0 |
| | 50 | No lesion | 0 |
| **ATV$^{WT}$:3D6** | 2 | No lesion | 0 |
| | 7 | No lesion | 0 |
| | 14 | No lesion | 0 |
| | 22 | No lesion | 0 |
| | 26 | No lesion | 0 |
| | 28 | No lesion | 0 |
| | 36 | No lesion | 0 |
| | 43 | No lesion | 0 |
| | 44 | No lesion | 0 |
| | 49 | No lesion | 0 |
| **ATV$^{cisLALA}$:3D6** | 5 | No lesion | 0 |
| | 10 | No lesion | 0 |
| | 11 | No lesion | 0 |
| | 24 | No lesion | 0 |
| | 29 | No lesion | 0 |
| | 32 | No lesion | 0 |
| | 40 | No lesion | 0 |
| | 41 | No lesion | 0 |
| | 46 | No lesion | 0 |
| | 51 | No lesion | 0 |

*The MRI score is a semi-quantitative assessment (value from 0 to 10) reflecting the extent of ARIA-like lesions for each animal. It takes into account the number of isolated lesions (from 1 lesion to 6 lesions), and the approximate size of all the lesions (from 1 to greater than 5) observed through the course of the study for a given animal. A number of lesions >5 indicates occurrence of multiple convexity lesions that cannot isolated and counted individually. Size of the lesions takes into account extent of a given lesion in a single scan and number of scans showing the lesion. A lesions size >5 (animal #15) reflects the presence of lesions in all scans. Both number and size of lesions are assessed blinded of the treatment allocation.
